# Noncanonical Rab9a action supports endosomal exit of human papillomavirus during virus entry

**DOI:** 10.1101/2023.05.01.538937

**Authors:** Jeongjoon Choi, Daniel DiMaio

## Abstract

Rab GTPases play key roles in controlling intracellular vesicular transport. GTP-bound Rab proteins support vesicle trafficking. Here, we report that, unlike cellular protein cargos, the delivery of human papillomaviruses (HPV) into the retrograde transport pathway during virus entry is inhibited by Rab9a in its GTP-bound form. Knockdown of Rab9a hampers HPV entry by regulating the HPV-retromer interaction and impairing retromer-mediated endosome-to-Golgi transport of the incoming virus, resulting in the accumulation of HPV in the endosome. Rab9a is in proximity to HPV as early as 3.5 h post-infection, prior to the Rab7-HPV interaction. HPV displays increased association with retromer in Rab9a knockdown cells, even in the presence of dominant negative Rab7. Thus, Rab9a can regulate HPV-retromer association independently of Rab7. Surprisingly, excess GTP-Rab9a impairs HPV entry, whereas excess GDP-Rab9a stimulates entry. These findings reveal that HPV employs a trafficking mechanism distinct from that used by cellular proteins.

## Introduction

Rab GTPases are key regulators of intracellular vesicular transport [1-3]. In general, when bound to guanosine triphosphate (GTP), Rab proteins are active and support vesicle trafficking, but they do not support vesicle transport when bound to guanosine diphosphate (GDP) [1, 3]. Trafficking of some viruses to the site of viral genome replication depends on cellular Rab proteins [4-6]. Human papillomaviruses (HPVs) are non-enveloped, double-stranded DNA viruses responsible for ∼5% of human cancer, including essentially all cervical cancer [7]. HPV entry requires various Rab proteins, but it remains largely unclear which Rab proteins are employed at which specific steps of virus entry [6]. Here, we show that a GDP-bound Rab protein is necessary for HPV trafficking from the endosome to the trans-Golgi network (TGN) during entry.

The HPV virion contains 360 molecules of the major capsid protein, L1, and up to 72 molecules of the minor capsid protein, L2 [8]. L1 is responsible for HPV binding to the cell surface, while L2 is essential for trafficking of the viral genome to the nucleus where viral gene expression and DNA replication occur [9]. Upon virus internalization, viral components including viral DNA reside within vesicular retrograde trafficking compartments throughout the entry process [10, 11]. After initially localizing in endosomes, HPV traffics to the *trans*-Golgi network (TGN) in a manner dependent on retromer, a cellular protein sorting complex [12, 13]. HPV also traffics through the Golgi apparatus and possibly the endoplasmic reticulum *en route* to the nucleus [9, 12-15]. During entry, a portion of the L2 protein protrudes through the endosome membrane into the cytoplasm, allowing it to interact with various cellular proteins to enable retrograde trafficking of the incoming virus particle [16, 17]. Cytoplasmic proteins involved in HPV retrograde trafficking that interact with L2 include retromer [13], sorting nexin 17 and 27 [18, 19], dynein [20], retriever [21] and COPI [22].

Retromer, a three-subunit protein complex (VPS26, VPS29, and VPS35), binds directly to the C-terminus of L2 and plays a critical role in endosome-to-TGN transport of HPV [12, 13]. GTP-Rab7 interacts with retromer [23] and recruits and stabilizes the association of HPV with retromer at the endosomal membrane [24]. In addition, cycling between GTP- and GDP-bound Rab7 is critical for endosome-to-TGN trafficking of HPV: while GTP-Rab7 recruits retromer, GTP hydrolysis to generate GDP-Rab7 is required for dissociation of the retromer from HPV, which must occur for trafficking to proceed [24]. Rab9a is also required for HPV entry, but its role in this process has not been studied [12, 15]. Because Rab9a facilitates endosome-to-TGN trafficking of cellular proteins [25], it is plausible that it acts at a similar step as Rab7 in HPV entry.

In this study, we show that Rab9a supports HPV type 16 (HPV16) entry by associating with incoming HPV early during infection and modulating the association of retromer and HPV. This activity of Rab9a is critical for endosome exit of HPV. Rab9a engages HPV prior to Rab7 and interferes with HPV-retromer association even in cells expressing dominant negative Rab7. Surprisingly, unlike cellular protein cargos, Rab9a-mediated HPV trafficking is promoted by an increased GDP-Rab9a to GTP-Rab9a ratio but inhibited by a decreased ratio of GTP-Rab9a to GDP-Rab9a. Our findings reveal a vesicle transport mechanism that requires GDP-bound Rab, distinct from cellular protein cargo trafficking.

## Results

### Rab9a is required for HPV trafficking from endosomes to Golgi during virus entry

Although Rab9a is required for efficient HPV pseudovirus (PsV) infection, its role in this process has not been investigated. To determine the role of Rab9a in HPV infection, HeLa S3 cells were transfected with <u>n</u>on-targeting <u>c</u>ontrol siRNA (siNC) and siRNA targeting Rab9a expression (siRab9a), and Rab9a knockdown was confirmed by Western blotting (Fig. 1A and S1A). These cells were then infected with HPV16 PsV consisting of a complete L1 and L2 capsid containing a reporter plasmid expressing GFP. A 3xFLAG tag was appended to the C-terminus of L2. Infectivity was determined by flow cytometry for GFP fluorescence at 48 <u>h p</u>ost-infection (hpi). This assay effectively measures only the entry phase of the HPV life cycle.

**Fig. 1.**
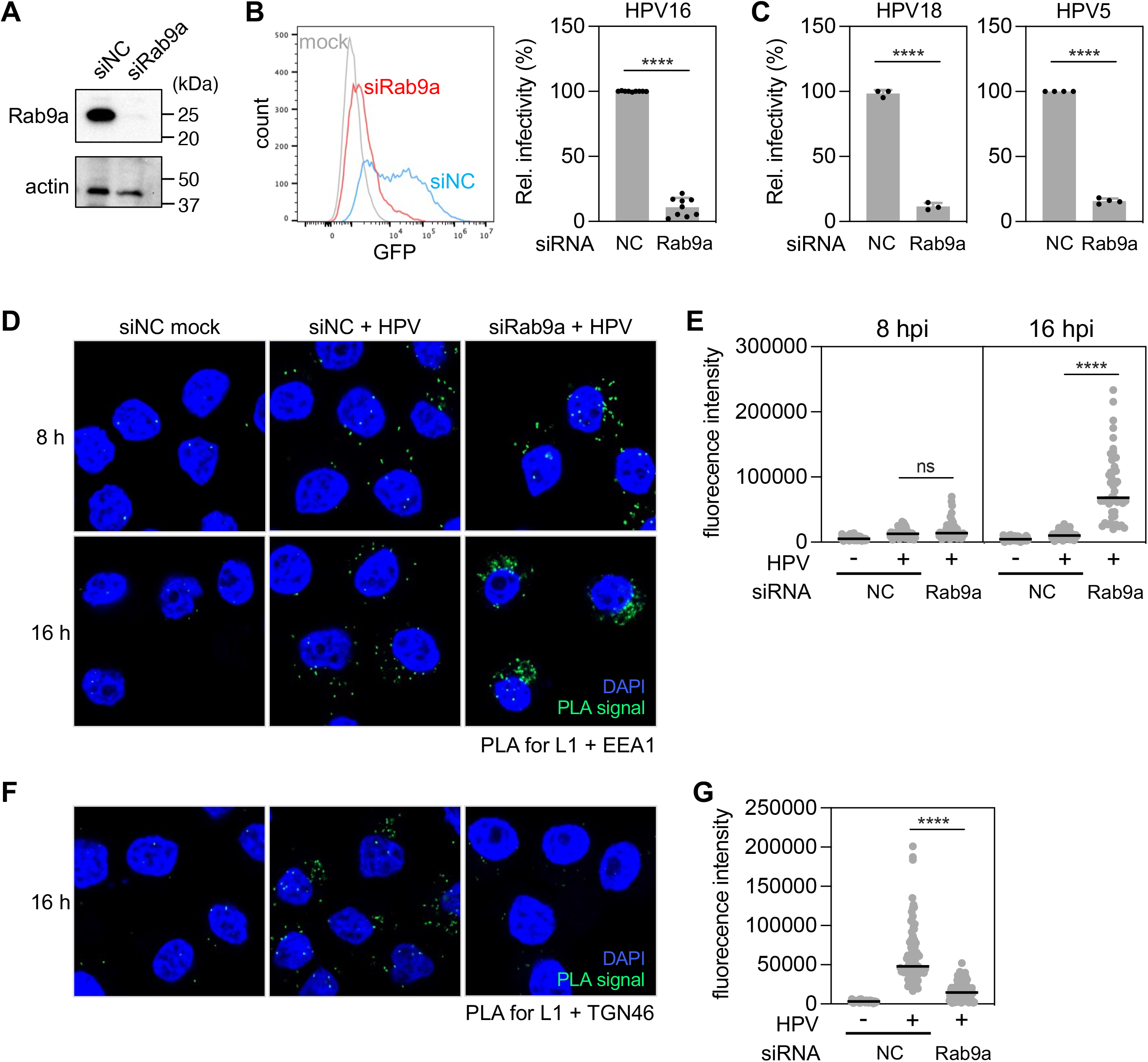
Rab9a knockdown inhibits HPV infection by impairing its trafficking from endosomes to Golgi. (**A**) HeLa S3 cells were transfected with negative control (siNC) or Rab9a-targeting siRNA (siRab9a) and subjected to Western blot analysis using an antibody recognizing Rab9a (top panel) and actin (bottom panel) as a loading control. (**B**) siRNA-treated cells as described in (**A**) were mock-infected or infected at the MOI of ∼2 with HPV16 PsV L2-3XFLAG containing the GFP reporter plasmid. At 48 hpi, GFP fluorescence was determined by flow cytometry. The results are shown as a histogram (*left*) and as percent relative infectivity (based on mean fluorescence intensity) normalized to the siNC treated cells (*right*). Each dot shows the result of an individual experiment. NC, siNC. ****, *P* < 0.0001. (**C**) As in (**B**) except cells were infected with HPV18 and HPV5. (**D**) HeLa S3 cells were transfected with siNC or siRab9a siRNAs and infected with HPV harboring the HcRed reporter plasmid at the MOI of ∼200. At 8 and 16 hpi, PLA was performed with antibodies recognizing HPV L1 and EEA1. Mock, uninfected; HPV, infected. PLA signals are green; nuclei are blue (DAPI). Similar results were obtained in two independent experiments. (**E**) The fluorescence of PLA signals was determined from multiple images obtained as in (**D**). Each dot represents an individual cell (*n*>40) and black horizontal lines indicate the mean value of the analyzed population in each group. ****, *P* < 0.0001; ns, not significant. The graph shows results of a representative experiment. Similar results were obtained in two independent experiments. (**F**) As in (**D**) except PLA was performed at 16 hpi with antibodies recognizing HPV L1 and TGN46. (**G**) Images as in (**F**) were analyzed as described in (**E**).

Consistent with our previously published results [12], knockdown of Rab9a inhibited HPV infectivity by ∼65-85% (Fig. 1B and S1B). Moreover, other pathogenic HPVs HPV18 and HPV5 also displayed reduced infectivity in HeLa cells depleted for Rab9a (Fig. 1C). Rab9a knockdown also inhibited HPV infection by ∼85% in human HaCaT skin keratinocytes (Fig. S1C and S1D). These data indicate that Rab9a is necessary for efficient HPV entry.

To determine the HPV entry step impaired by Rab9a knockdown, we used proximity ligation assay (PLA) to examine HPV localization in cells infected with HPV16 PsV. PLA generates a fluorescent signal in intact cells when two proteins of interest are within 40 nm [26]. Because Rab9a is necessary for cellular protein trafficking from the endosome to Golgi [25], we examined the proximity of HPV L1 and marker proteins for the endosome (EEA1) or TGN (TGN46). There were negligible PLA signals in mock infected cells. At 8 hpi, similar levels of L1-EEA1 PLA signals were observed in cell transfected with either siNC or siRab9a (Fig. 1D and 1E), indicating that Rab9a is not required for HPV endocytosis. In contrast, at 16 hpi, a time when most HPV has largely exited the endosome and entered the TGN in normal cells [13, 14], Rab9a knockdown resulted in much stronger L1-EEA1 PLA signals than in the control cells, indicating that HPV accumulates in endosomes in the absence of Rab9a (Fig. 1D and 1E). Consistent with this result at 16 hpi, control infected cells displayed strong L1-TGN46 PLA signals, as expected, whereas Rab9a knockdown cells showed very little signal (Fig. 1F and 1G), indicating that HPV arrival in TGN was impaired in the absence of Rab9a. These data indicate that Rab9a supports HPV entry by enabling trafficking of HPV from the endosome to the TGN.

### Rab9a knockdown regulates HPV-retromer interaction

Knockdown of retromer subunit VPS35 or VPS29 or infection with a PsV harboring retromer binding site mutations in L2 cause HPV accumulation in endosomes, indicating that the HPV-retromer interaction is required for exiting endosomes (Fig. S2) [13, 16]. Since these phenotypes are similar to that caused by Rab9a knockdown described above, we used PLA to test whether Rab9a knockdown affected HPV interaction with retromer. Figures 2A and 2B show that the L1-VPS35 interaction as measured by PLA at 8 and 16 hpi was increased by Rab9a knockdown. These data suggest that Rab9a facilitates endosome exit by regulating the HPV-retromer interaction.

**Fig. 2.**
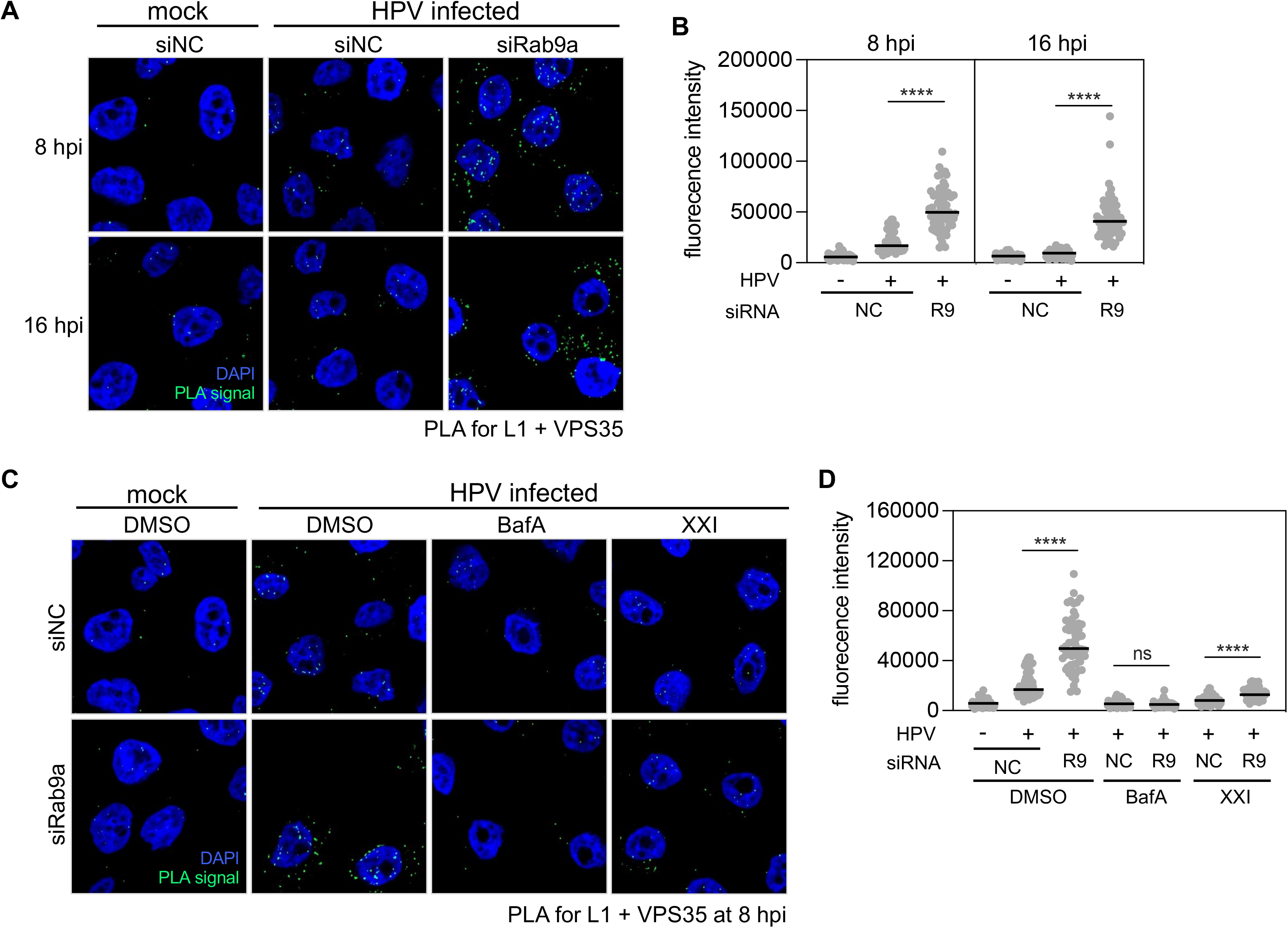
Rab9a knockdown increases HPV-retromer association. (**A**) HeLa S3 cells were transfected with siNC or siRab9a and infected at the MOI of ∼200 with HPV16 PsV L2-3XFLAG containing the HcRed reporter plasmid. At 8 or 16 hpi, PLA was performed with antibodies recognizing HPV L1 and VPS35. Mock, uninfected. PLA signals are green; nuclei are blue (DAPI). Similar results were obtained in three independent experiments. (**B**) The fluorescence of PLA signals was determined from multiple images obtained as in (**A**). Each dot represents an individual cell (*n*>40) and black horizontal lines indicate the mean value of the analyzed population in each group. NC, siNC; R9, siRab9a. ****, *P* < 0.0001; ns, not significant. The graph shows results of a representative experiment. Similar results were obtained in three independent experiments. (**C**) As in (**A**) at 8 hpi except DMSO, Bafilomycin A1 (BafA), or γ-secretase inhibitor (XXI) were added to the medium 30 min prior to infection. (**D**) Images as in (**C**) were analyzed as described in (**B**). The graph shows results of a representative experiment. Similar results were obtained in two independent experiments.

Endosomal acidification and γ-secretase are required for HPV entry and retromer association at relatively early times during infection by supporting L2 insertion into the endosome membrane [27, 28]. γ-secretase may also act at a later step of infection [29]. When endosomal acidification was inhibited by Bafilomycin A1 (BafA), the HPV-retromer interaction at 8 hpi was abolished in the presence or absence of Rab9a (Fig. 2C and 2D). Similarly, inhibition of γ-secretase by compound XXI also greatly reduced HPV-retromer association and largely eliminated the increase in the HPV-retromer interaction caused by Rab9a knockdown (Fig. 2C and 2D). Collectively, these data show that endosomal acidification and γ-secretase activity are required for both HPV-retromer interaction in cells with unperturbed Rab9a expression and the increased HPV-retromer association caused by Rab9a knockdown, implying that acidification and γ-secretase are required prior to Rab9a action during entry.

### Rab9a is in close proximity to HPV at early times post-infection

We used PLA to test whether HPV and Rab9a are in proximity during HPV entry. As shown in Fig. 3A and 3B, there were clear L1-Rab9a PLA signals at 8 hpi, showing that L1 and Rab9a are in close proximity at this time. To determine when Rab9a associates with HPV during entry, we conducted L1-Rab9a PLA at various times after infection. This analysis revealed that Rab9a is in proximity to L1 as early as 3.5 hpi (Fig. 3A and 3B), showing that HPV associates with Rab9a relatively early during virus entry.

**Fig. 3.**
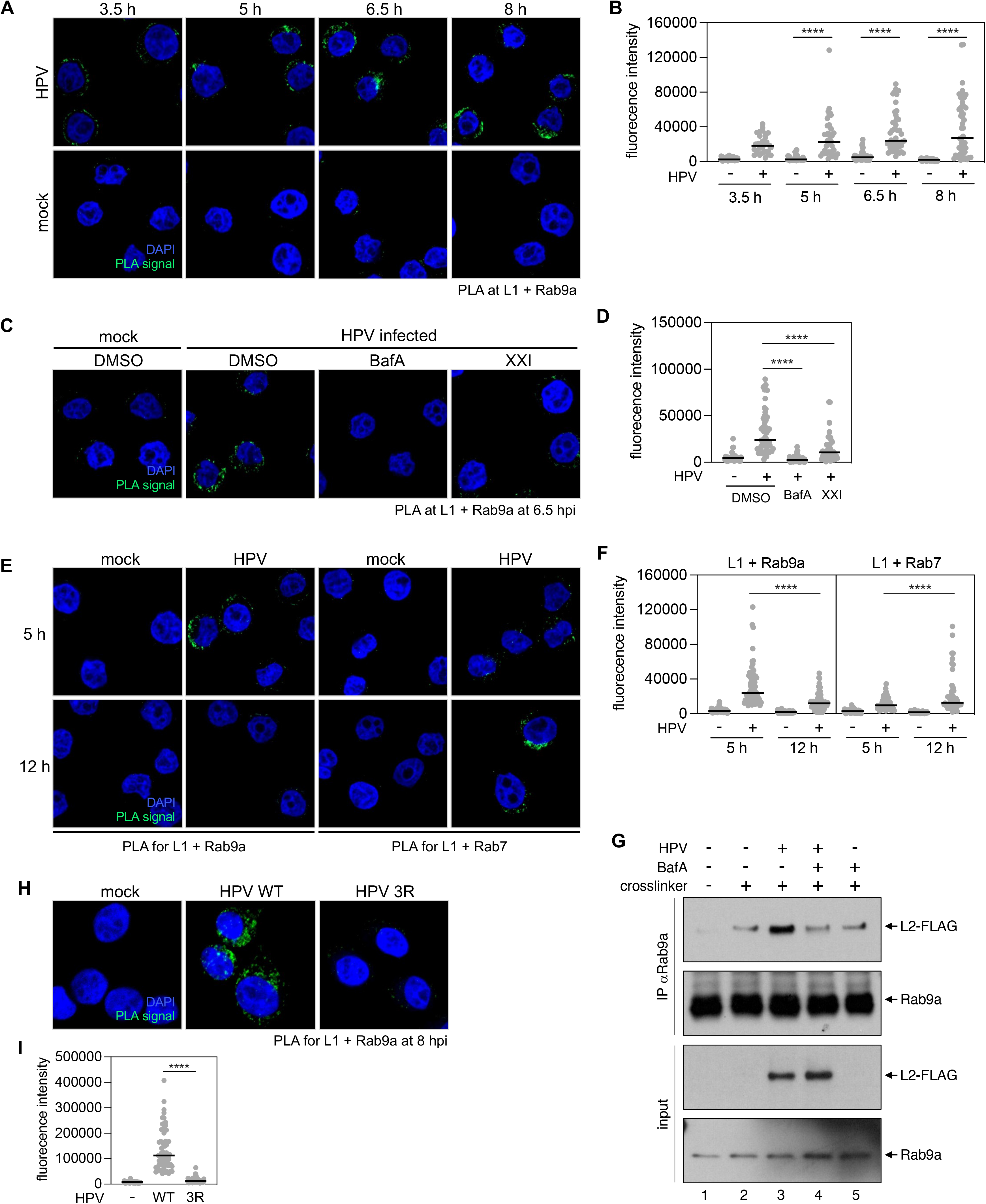
Rab9a is in proximity to HPV at early times post-infection prior to Rab7. (**A**) HeLa S3 cells were mock-infected or infected at the MOI of ∼200 with HPV16 PsV L2-3XFLAG containing the HcRed reporter plasmid. At 3.5, 5, 6.5, or 8 hpi, PLA was performed with antibodies recognizing HPV L1 and Rab9a. PLA signals are green; nuclei are blue (DAPI). Similar results were obtained in two independent experiments. (**B**)The fluorescence of PLA signals was determined from multiple images obtained as in (**A**). Each dot represents an individual cell (*n*>30) and black horizontal lines indicate the mean value of the analyzed population in each group. ****, *P* < 0.0001. The graph shows results of a representative experiment. Similar results were obtained in two independent experiments. (**C**) As in (**A**) at 6.5 hpi except DMSO, Bafilomycin A1 (BafA), or γ-secretase inhibitor (XXI) were added to the medium 30 min prior to infection. Similar results were obtained in two independent experiments. (**D**) Images as in (**C**) were analyzed as described in (**B**). (**E**) As in (**A**) except PLA was performed at 5 and 12 hpi with antibodies recognizing HPV L1 and Rab9a or Rab7. Similar results were obtained in two independent experiments. (**F**) Images as in (**E**) were analyzed as described in (**B**). (**G**) HeLa S3 cells infected at the MOI of ∼100 with HPV16 PsV L2-3XFLAG (HPV) harboring the HcRed reporter plasmid. DMSO (-) or Bafilomycin A1 (BafA, +) were added to the medium 30 min prior to infection. DSP crosslinker was added as indicated. At 8 hpi, lysates were pulled down with antibody recognizing Rab9a (IP) and subjected to Western blot analysis using antibodies recognizing FLAG, or Rab9a, together with samples not pulled down (input). (**H**) As in (**A**) at 8 hpi except HPV16 PsV containing HA-tagged wild-type (WT) and 3R mutant L2 were used. Similar results were obtained in two independent experiments. (**I**) Images as in (**H**) were analyzed as described in (**B**).

We then used PLA to test whether endosome acidification and γ-secretase activity are required for Rab9a association with HPV. As shown in Fig. 3C and 3D, at 6.5 hpi, inhibition of endosomal acidification or γ-secretase activity abrogated or reduced L1-Rab9a association, consistent with our findings that acidification and γ-secretase are required upstream of Rab9a during infection (Fig. 2C and 2D).

Rab7, which is also required for HPV entry [24], is found primarily in late endosomes [25, 30], although there is evidence that Rab7 is also associated with early endosomes [31, 32]. To test whether HPV associates with Rab9a prior to its association with Rab7, we performed PLA for L1-Rab9a and L1-Rab7 at 5 and 12 hpi. PLA signals from L1-Rab9a were high at 5 hpi and decreased at 12 hpi, whereas L1-Rab7 PLA signals were low at 5 hpi and increased at 12 hpi (Fig. 3E and 3F). These results indicate that HPV comes in proximity to Rab9a prior to Rab7. Taken together, these data indicate that Rab9a localizes in proximity to HPV at early times during infection prior to Rab7 association in a manner dependent on endosome acidification and γ-secretase action.

### Rab9a associates with L2 upon protrusion through the endosomal membrane

L2 interacts with several cytoplasmic proteins involved in HPV retrograde trafficking [13, 18-22]. To test if Rab9a is in a complex with L2, we performed immunoprecipitation experiments in cells infected with PsV containing FLAG-tagged L2. At 6.5 hpi, cells were treated with the cell-permeable cross-linker dithiobis(succinimidyl propionate) (DSP) for 30 min and lysed in buffer containing detergents (1% *n*-Dodecyl β-D-maltoside [DDM] and 0.05% Triton X-100). Anti-Rab9a antibody was then used to immunoprecipitate Rab9a and associated proteins. The immunoprecipitates were subjected to gel electrophoresis and probed with anti-FLAG antibody. Although there is a faint background band that comigrates with L2 in uninfected cells treated with the cross-linker (Fig. 3G, lane 2), anti-Rab9a co-immunoprecipitated the L2 protein from infected cells (Fig. 3G, lanes 2 and 3), indicating that L2 and Rab9a are present in a protein complex. Similar to L1-Rab9a association determined by PLA (Fig. 3E and 3F), inhibition of the endosomal acidification with BafA1 abolished the L2-Rab9a interaction as assessed by co-immunoprecipitation (Fig. 3G, lanes 3, 4 and 5). As endosome acidification is also required for L2 protrusion through the endosome membrane [16, 27, 28], L2 protrusion may be required for Rab9a-HPV association. To test this, cells were infected with an L2 mutant in which the cell-penetrating peptide sequence RKRRKR was replaced by RRR (this 3R mutant is impaired for membrane insertion and cytoplasmic protrusion [16, 33]), and L1-Rab9a association was assessed at 8 hpi by PLA. The 3R mutant abolished the L1-Rab9a association, unlike wild-type (Fig. 3H and 3I), supporting the conclusion that the L2 protrusion is necessary for HPV-Rab9a association.

### Rab9a knockdown increases the abundance of GTP-bound Rab7

Because knockdown of either Rab9a or Rab7 affects HPV-retromer association, we wondered whether Rab9a affects Rab7 activity. To test whether Rab9a knockdown affected the abundance of GTP-bound Rab7, we performed pull-downs with Rab-interacting lysosomal protein (RILP), which binds GTP-bound but not GDP-bound Rab7 [34]. Purified glutathione S-transferase (GST) or the GST-RILP fusion proteins were incubated with extracts from uninfected HeLa cells transfected with various siRNAs and then immunoprecipitated with anti-GST antibody. Rab7 was co-immunoprecipitated from control cells transfected with siNC incubated with GST-RILP but not from cells incubated with the negative control GST (Fig. 4A, lanes 1 and 2). Knockdown of the Rab7 GTPase-activating protein (GAP) TBC1D5 caused about 2.5-fold increase in GTP-Rab7, as expected (Fig. 4A, compare lanes 2 and 3). Approximately 4-fold more Rab7 was pulled down from extracts from the cells transfected with siRab9a than from the control cells (Fig. 4A, compare lanes 2 and 4) while total Rab7 abundance was unaltered (Fig. 4A, second panel from top). This result indicates that Rab9a knockdown increases amount of GTP-Rab7. Moreover, Rab9a knockdown increased the localization of Rab7 to the endosome as shown by increased Rab7-EEA1 PLA signal (Fig. 4B and 4C) and increased Rab7-VPS35 association at 12 hpi in infected cells as measured by PLA (Fig. 4D and 4E), consistent with the ability of GTP-Rab7 – not GDP-Rab7 – to bind the retromer complex [35]. The increase of Rab7-retromer interaction occurred both in the absence of HPV infection and to a greater extent in cells infected with HPV (Fig. 4D and 4E). We previously used co-immunoprecipitation to show that Rab7 and HPV are in a complex at 12 hpi [24]. Here, we used PLA to demonstrate that the L1-Rab7 association in HPV infected cells was increased by Rab9a knockdown as early as 8 hpi (Fig. S3A and S3B). Taken together, these data indicate that Rab9a knockdown promotes an increase of GTP-Rab7, the canonical activated form of Rab7.

**Fig. 4.**
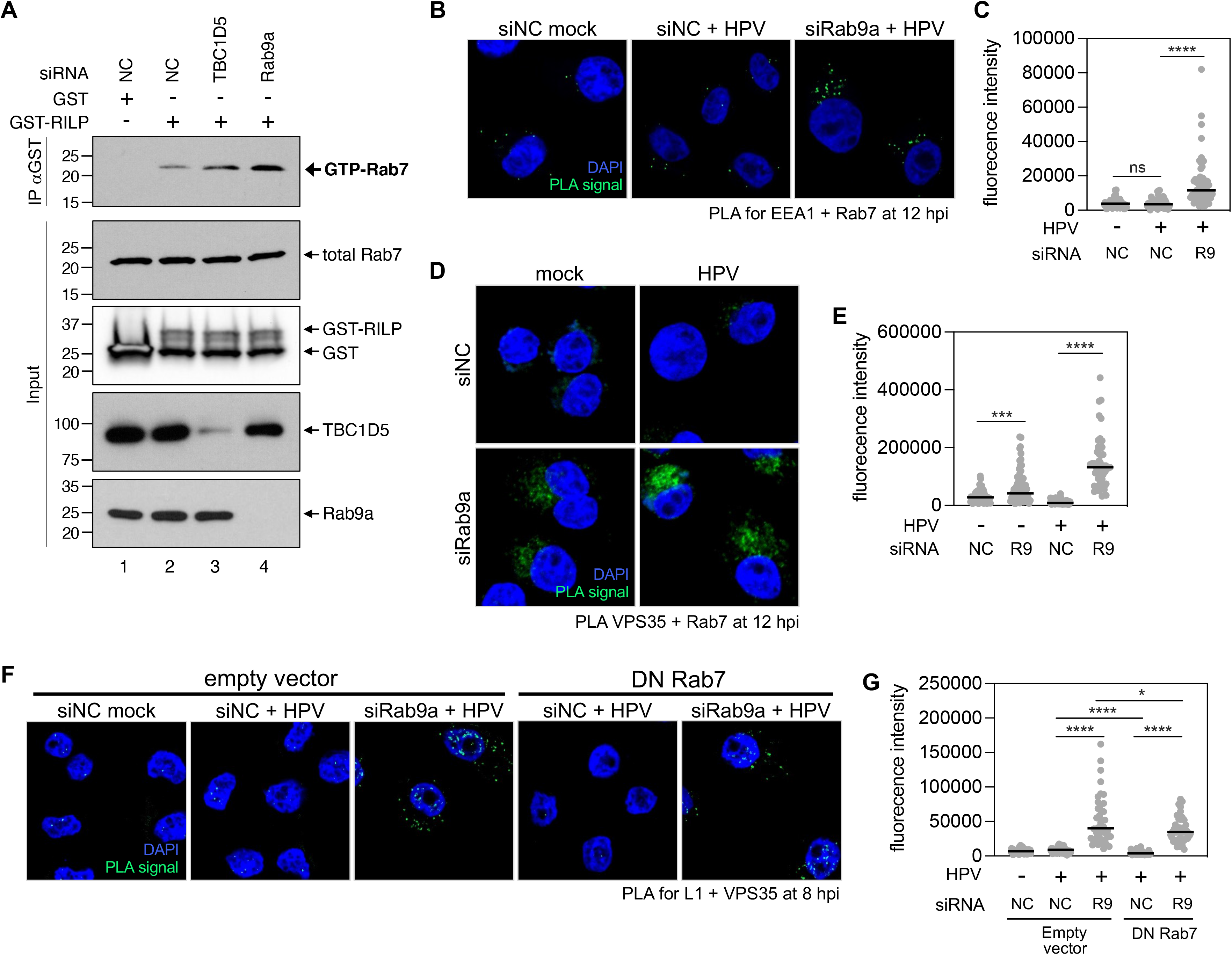
Rab9a knockdown enhances HPV-retromer association even in cells overexpressing GDP-Rab7 despite increases GTP-Rab7 abundance. (**A**) HeLa S3 cells were transfected with siNC, siRab9a or siTBC1D5. At 48 h after transfection, lysates were pulled down with GST or GST-RILP (IP) and subjected to Western blot analysis together with samples not pulled down (input) using antibodies recognizing Rab7, GST, TBC1D5, or Rab9a. Similar results were obtained in two independent experiments. (**B**) HeLa S3 cells were transfected with siNC or siRab9a and mock-infected or infected at the MOI of ∼200 with HPV16 PsV L2-3XFLAG containing the HcRed reporter plasmid. At 12 hpi, PLA was performed with antibodies recognizing EEA1 and Rab7. PLA signals are green; nuclei are blue (DAPI). Similar results were obtained in two independent experiments. (**C**) The fluorescence of PLA signals was determined from multiple images obtained as in (**B**). Each dot represents an individual cell (*n*>30) and black horizontal lines indicate the mean value of the analyzed population in each group. ***, *P* < 0.001; ****, *P* < 0.0001. The graph shows results of a representative experiment. Similar results were obtained in two independent experiments. (**D**) As in (**B**) except PLA was performed at 12 hpi with antibodies recognizing VPS35 and Rab7. Similar results were obtained in two independent experiments. (**E**) Images as in (**D**) were analyzed as described in (**C**). (**F**) As in (**B**) except HeLa S3 cells stably transduced with an empty vector or a plasmid expressing dominant negative Rab7 (Rab7 DN) were used and PLA was performed at 8 hpi with antibodies recognizing HPV L1 and VPS35. Similar results were obtained in two independent experiments. (**G**) Images as in (**F**) were analyzed as described in (**C**). *, *P* < 0.05.

### Rab9a knockdown increases HPV-retromer association even in the presence of dominant negative Rab7

GTP-bound Rab7 is critical for retromer recruitment to endosomes [35]. Indeed, as reported previously, cells expressing a dominant negative Rab7 mutant (T22N; DN Rab7), which binds only GDP and is thus unable to interact with retromer [23, 35], displayed reduced L1-VPS35 PLA signals at 8 and 16 hpi in control cells expressing endogenous Rab9a (Fig. 4F, 4G, and S4) [23, 24], showing that GTP-Rab7 supports HPV-retromer association. To test whether Rab9a regulates HPV-retromer association by decreasing GTP-Rab7, we analyzed cells expressing DN Rab7, which contain approximately only 5% GTP-Rab7 compared to control cells (Fig. S5). As shown in Fig. 4F, 4G, and S4, the L1-VPS35 PLA signals at 8 and 16 hpi in Rab9a knockdown cells were elevated to a similar level in the presence and absence of DN Rab7. Thus, when Rab9a is knocked down, GTP-Rab7 is not strictly required for retromer-HPV association. These results suggest that Rab9a regulates retromer association independently of Rab7, consistent with the finding that HPV associates with Rab9 before it associates with Rab7.

### Excess GTP-bound Rab9a inhibits HPV infection whereas excess GDP-bound Rab9a promotes it

The GTP-bound form of Rab proteins typically promotes cargo trafficking [1]. To determine which form of Rab9a (Fig. 5A) is necessary for HPV infection, we transiently introduced constitutively active (CA) HA-tagged Rab9a mutant (Q66L) into 293TT cells. Cells expressing CA Rab9a likely accumulate GTP-Rab9a and have less GDP-Rab9a, compared to normal cells (Fig. 5A and 5B). 48 h after transfection, the cells were infected with HPV16 PsV. Forty-eight hpi we used flow cytometry to quantify GFP fluorescence as a measure of infection and anti-HA staining to document successful expression of the mutant Rab9a protein. This experimental design allows us to compare HPV infection in the presence or absence of CA Rab9a (i.e., in HA-positive and HA-negative cells, respectively) in the same set of infected cell population (Fig. 5C). Surprisingly, unlike cellular cargo, cells expressing CA Rab9a with presumably an increased GTP-Rab9a to GDP-Rab9a ratio reduced HPV infection (Fig. 5C), indicating that excess GTP-Rab9a impairs HPV infection.

**Fig. 5.**
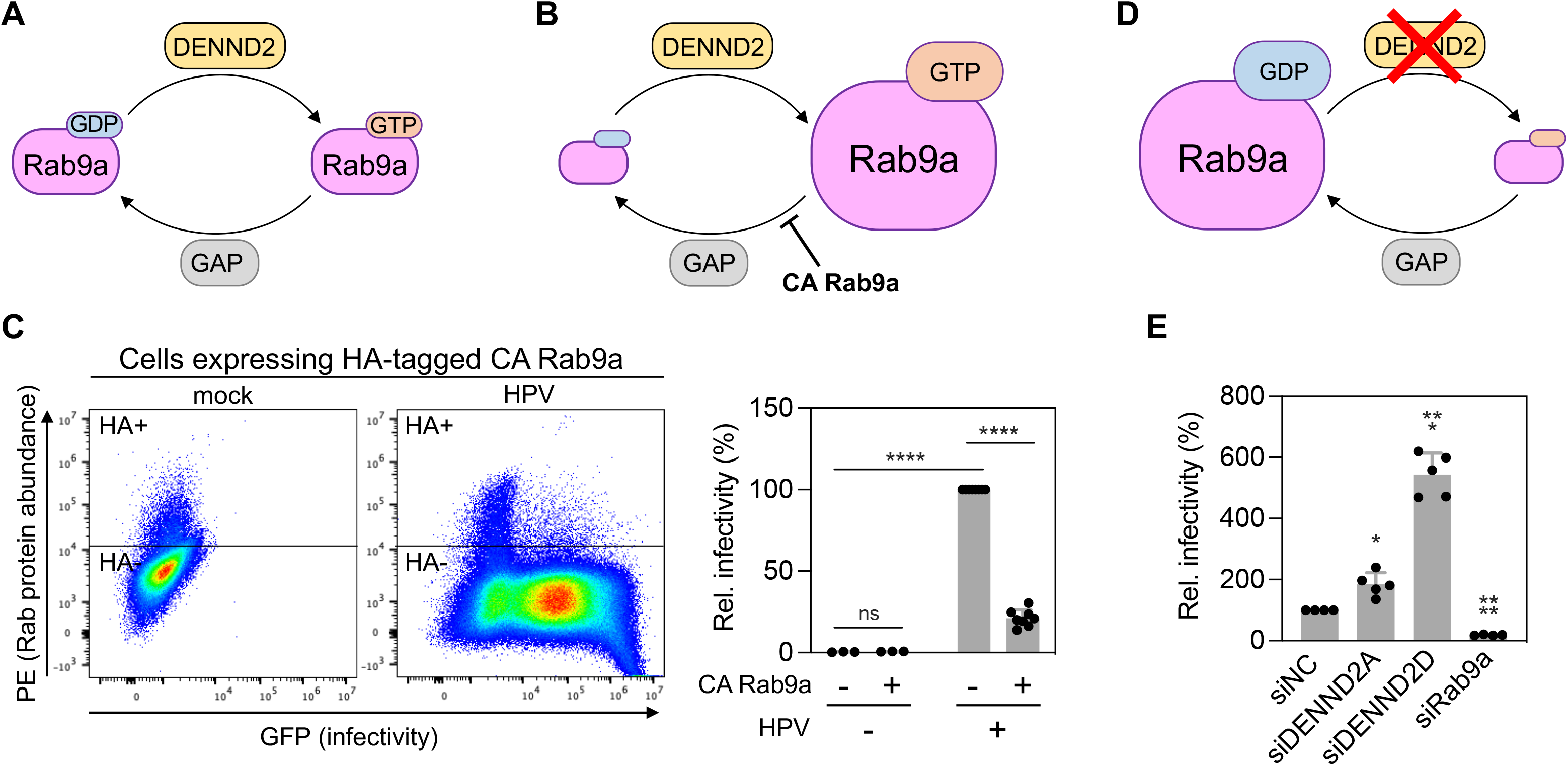
GTP-bound Rab9a inhibits HPV infection but GDP-bound Rab9a promotes it. (**A**) Schematic of Rab9a cycling between GTP-bound and GDP-bound forms. DENND2, a Rab9a GEF, converts GDP-Rab9a to GTP-Rab9a by exchanging GDP to GTP. A GTPase-activating protein (GAP) hydrolyzing GTP-Rab9a is yet to be identified. (**B**) Constitutively active (CA) Rab9a mutant is locked in GTP-bound form; thus cells expressing CA Rab9a accumulate GTP-Rab9a. (**C**) 293TT cells were transfected with a plasmid expressing HA-tagged CA Rab9a. At 48 h after transfection, cells were infected at the MOI of ∼2 with HPV16 PsV L2-3XFLAG containing the GFP reporter plasmid. At 48 hpi, samples were stained using PE-conjugated antibody recognizing HA, and flow cytometry was used to determine PE and GFP fluorescence. Cells expressing HA-tagged CA Rab9a were marked as HA+, and those not expressing such proteins were marked as HA-. Representative dot blots are shown in left panels. On right, data from multiple repeats are shown as percent relative infectivity normalized to populations not expressing Rab9a CA (set at 100%). Each dot shows the result of an individual experiment. ****, *P* < 0.0001; ns, not significant. (**D**) Knockdown of DENND2 inhibits the conversion of GDP-Rab9a to GTP-Rab9a, thereby accumulating GDP-Rab9a. (**E**) HeLa S3 cells transfected with siNC, siDENND2A, siDENND2D, or siRab9a were infected at the MOI of ∼0.2-0.4 with HPV16 PsV L2-3XFLAG (HPV) harboring the GFP reporter plasmid. At 48 hpi, GFP fluorescence was determined by flow cytometry. The results were shown in percent relative infectivity normalized to the siNC treated cells (set at 100%). Each dot shows the result of an individual experiment. *, *P* < 0.05; ***, *P* < 0.001; ****, *P* < 0.0001.

Next, we determined the effect of excess GDP-Rab9a on HPV infection. We could not test DN Rab9a mutant (S21N) using the approach described above because we were unable to generate a large enough number of cells expressing DN Rab9a. As an alternative approach to generate cells with excess GDP-Rab9a, we knocked down two isoforms of DENND2 (a guanine nucleotide exchange factor (GEF) for Rab9a), which should cause accumulation of GDP-bound Rab9a (Fig. 5D). DENND2D knockdown resulted in 5-6-fold higher HPV infectivity compared to cells transfected with the control siRNA (Fig. 5E), while DENND2A knockdown resulted in ∼2-fold higher infectivity (Fig. 5E). This difference is consistent with the higher Rab9a GDP-exchanging activity of DENND2D compared to DENND2A [36]. The DENND2D knockdown also increased HPV infection in HaCaT cells, as in HeLa cells (Fig. S6A and S6B). Similarly, HPV18 and HPV5 also showed increased infection in DENND2D knockdown cells (Fig. S6C). Thus, HPV infection was inhibited by GTP-bound CA Rab9a and stimulated by excess GDP-Rab9a caused by knockdown of DENND2, a Rab9a GEF. Collectively, these data indicate that an increased ratio of GDP-Rab9a to GTP-Rab9a promotes HPV entry, whereas an increased ratio of GTP-Rab9a to GDP-Rab9a impairs it.

### Cellular cargo trafficking is promoted by GTP-Rab9a

Rab9a is required for the retrograde trafficking of cellular proteins, such as cation-independent mannose-6-phosphate receptors (CI-MPR) [25]. Unlike HPV trafficking as described above, endosome to TGN transport of CI-MPR is impaired by the increase of GDP-Rab9a in cells expressing Rab9a DN or depleted of DENND2 [36, 37], indicating that CI-MRP trafficking requires GTP-Rab9a, as expected. To investigate further the role of GTP-Rab9a action in cellular protein cargo trafficking in our cell system, we used immunofluorescence to monitor the effect of modulating Rab9a on another cellular retromer cargo protein, divalent metal transporter 1 isoform II (DMT1-II). Consistent with previously reported phenotypes on CI-MPR trafficking [36, 37], CA Rab9a increased the co-localization of DMT1-II and TGN46 in 293TT cells (Fig. 6A and 6B) whereas DENND2D knockdown cells showed decreased co-localization of DMT1-II and TGN46 compared to the control cells (Fig. 6C and 6D). These results suggest that, in contrast to HPV trafficking, increased GTP-Rab9a to GDP-Rab9a ratio promotes trafficking of DMT1-II as well as CI-MPR from endosome to TGN. These data indicate that Rab9a acts differently on cellular protein transport and HPV trafficking (Fig. 7A).

**Fig. 6.**
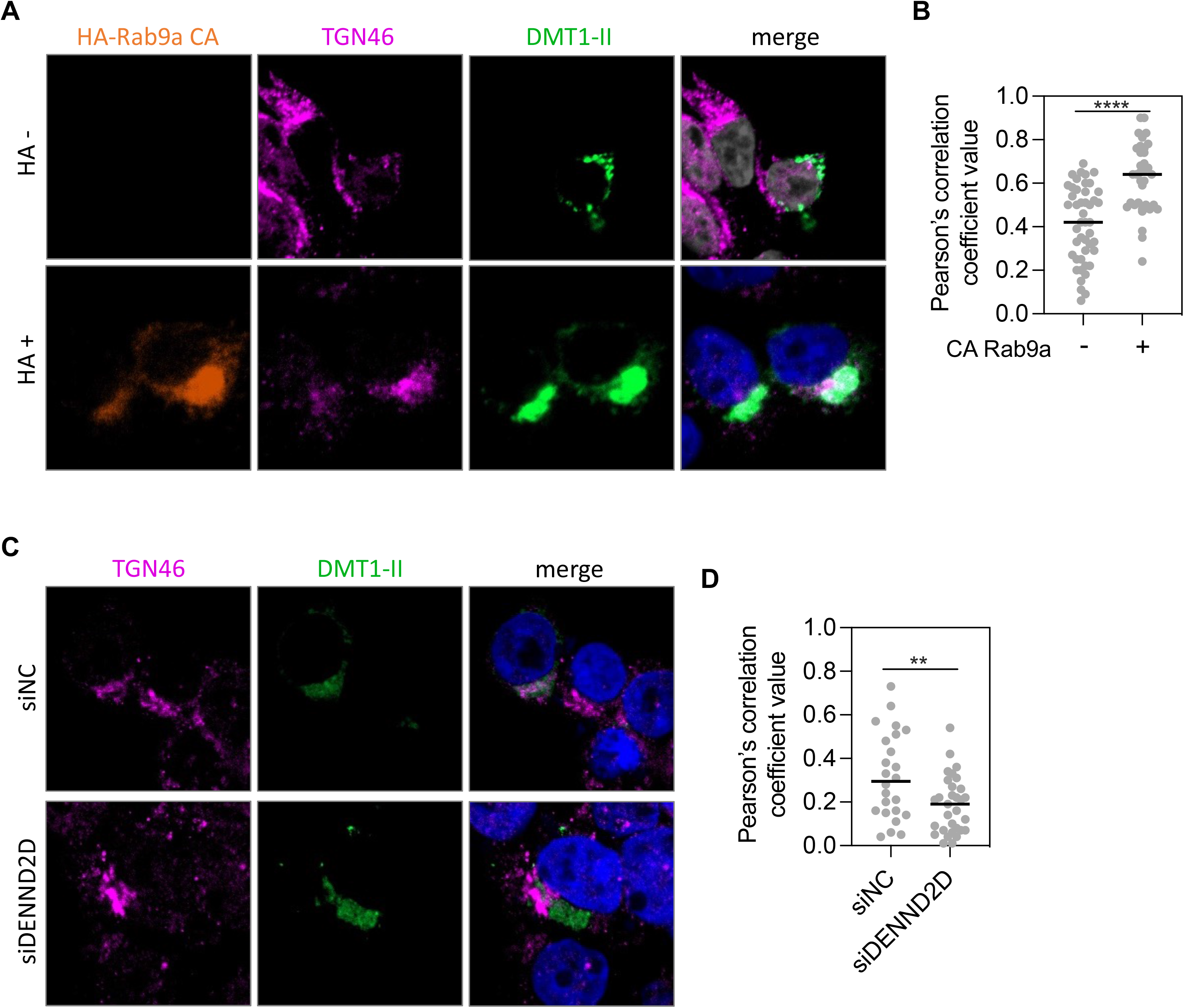
GTP-Rab9a promotes trafficking of DMT1-II. (**A**) 293TT cells were transfected with a plasmid expressing HA-Rab9a CA, followed by transfection of a DMT1-II expressing plasmid 24 h after the first transfection. Expression of HA-Rab9a CA was determined by anti-HA staining, which distinguishes cells expressing HA-Rab9a CA (+) or those not expressing it (-). DMT1-II and TGN46 were stained using antibodies recognizing GFP and TGN46, respectively. Immunofluorescence (IF) images were shown; HA-Rab9a CA, orange; TGN46, magenta; DMT1-II, green; nuclei (DAPI), blue. Merged image shows TGN46 and DMT1-II with overlap colored white. Similar results were obtained in three independent experiments. (**B**) Pearson’s correlation coefficient values for TGN46 and DMT1-II colocalization in those cells are shown. Each dot represents an individual cell (*n*>25) and black horizontal lines indicate the mean value of the analyzed population in each group. ****, *P* < 0.0001. The graph shows results of a representative experiment. Similar results were obtained in three independent experiments. (**C**) cells were transfected with siNC or siDENND2D, followed by transfection of a DMT1-II expressing plasmid 48 h after the first transfection and stained for TGN46 and DMT1-II as in (**A**). Similar results were obtained in two independent experiments. (**D**) Images as in (**C**) were analyzed as described in (**B**). **, *P* < 0.01

**Fig. 7.**
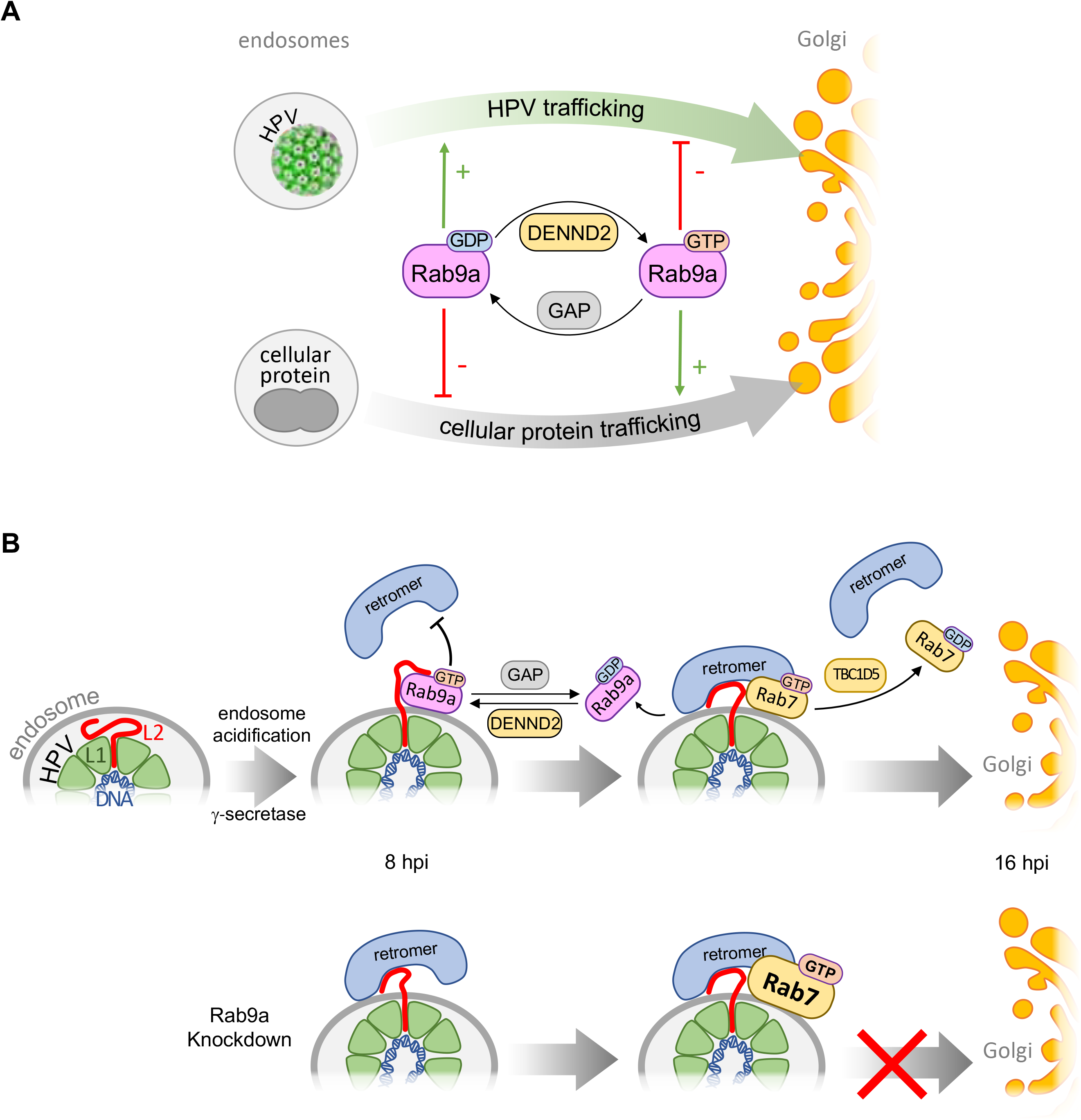
GDP-bound Rab9a promotes HPV entry by favoring the (retromer-mediated) retrograde HPV trafficking. (**A**) Retrograde transport of cellular protein(s), such as CI-MPR and DMT1-II, from endosome to Golgi requires the GTP-bound Rab9a and is impaired by GDP-bound Rab9a, as is the case for other known Rab proteins. Surprisingly, however, HPV entry is stimulated by GDP-bound Rab9a, previously thought to be an inactive form, whereas the GTP-bound Rab9a inhibits the virus entry. DENND2 impairs HPV entry by decreasing GDP-Rab9a and increasing GTP-Rab9a. (**B**) (*top*) Internalized HPV viral components including viral DNA reside within endosomes. Upon endosomal acidification and γ-secretase action, the L2 protein protrudes through the endosome membrane into the cytoplasm, allowing it to interact with various cellular proteins, including retromer, to facilitate retrograde trafficking of the incoming virus particle. Rab9a localizes in proximity to HPV early during infection, interacts with the HPV L2 protein, and impairs the HPV-retromer association at early times post-infection, prior to Rab7 localization to the HPV-containing vesicle. Thus, this early action of Rab9a appears to be independent of the GTP-bound Rab7. Later times post-infection, Rab7 localizes in proximity to HPV while Rab9a in proximity to HPV decreases. Hydrolysis of GTP-bound Rab7 causes dissociation of retromer, allowing HPV trafficking towards Golgi (*trans*-Golgi). (*bottom*) Rab9a knockdown increases retromer-HPV association at 8 hpi, which appears to be independent of Rab7. At later times, Rab9a knockdown results in increased GTP-bound Rab7 amounts, further enhancing retromer association, and thereby impairing the endosomal exit of HPV.

## Discussion

Rab GTPases play key regulatory roles in intracellular vesicular trafficking [1, 2, 4-6]. Like other small GTPases, Rab proteins exist in two forms, an active GTP-bound state that is necessary for trafficking of cellular cargo and an inactive GDP-bound state [1, 3]. Consistent with this view, excess GTP-Rab9a supports the transport of cellular cargo, such as CI-MPR and DMT1-II (Fig. 6) [36, 37]. Surprisingly, however, we show that HPV entry is impaired by excess GTP-Rab9a and stimulated by GDP-Rab9a. These findings indicate that HPV utilizes a trafficking mechanism different from that used by the cellular protein cargos (Fig. 7A).

Retrograde trafficking of HPV from endosome to TGN requires the interaction of retromer with L2 at vesicles containing HPV [12, 13]. The findings reported here as well as our previously published work suggest that the association of retromer to HPV-containing vesicles is regulated by different Rab proteins at different times post infection (Fig. 2 and 7B). We previously reported that GTP-Rab7 recruits retromer to HPV at the endosome membrane during HPV entry and that TBC1D5-catalyzed hydrolysis of GTP-Rab7 to GDP-Rab7 is required for dissociation of HPV from retromer, a step required for trafficking to proceed [1, 3, 6, 12, 15]. Thus, both GTP-Rab7 and GDP-Rab7 are required for HPV infection. In contrast, the GTP- and GDP-bound forms of Rab9a have opposing effects on HPV infection, with excess GDP-Rab9a supporting infection and excess GTP-Rab9a inhibiting it. Although both Rab9a knockdown and excess GTP-Rab9a impair HPV infection, the mechanism of inhibition is likely different: Rab9a knockdown depletes both forms of Rab9a (i.e., GTP-Rab9a and GDP-Rab9a) and increases HPV-retromer association, whereas excess GTP-Rab9a (i.e., cells expressing CA Rab9a) probably interferes with HPV-retromer association.

A previous study demonstrated that Rab9a co-immunoprecipitated with VPS35 [38], raising the possibility that Rab9a may regulate the association of retromer to the HPV-containing vesicle by interacting with the retromer complex. In support of this possibility, Rab9a associates with HPV during infection prior to the HPV association with Rab7, and Rab9a knockdown increased HPV-retromer association at 8 and 16 hpi. Alternatively, as both Rab9a and Rab7 interact with retromer [35, 38], Rab9a may impede the ability of Rab7 to bind retromer, thereby delaying the Rab7-retromer association until the proper time. Indeed, Rab9a knockdown increased Rab7 association with retromer at 12 hpi. Rab9a knockdown also increased HPV-retromer association at 8 hpi, whereas CA Rab7 did not increase HPV-retromer association at this relatively early time [24]. It is not clear why Rab9a knockdown increases the amount of GTP-Rab7.

Endosomal accumulation of HPV in Rab9a knockdown cells is likely a downstream consequence of increased HPV-retromer association. Consistent with this idea, Rab9a knockdown elevated HPV-retromer association at both 8 and 16 hpi, whereas HPV accumulation in endosomes was observed at 16 hpi - but not at 8 hpi - in Rab9a knockdown cells. How then might Rab9a depletion cause HPV endosomal accumulation? One possibility is that the increased GTP-Rab7 abundance in Rab9a knockdown cells prevents HPV-retromer dissociation, consistent with our previous findings that excess GTP-Rab7 due to expression of CA Rab7 or depletion of TBC1D5 increase HPV-retromer association and cause HPV accumulation in endosomes [24, 34, 39]. However, Rab9a knockdown increases retromer-HPV interaction even in cells expressing DN Rab7 (Fig. 4A and 4B), whereas cells expressing DN Rab7 in the absence of Rab9a knockdown fail to support the HPV-retromer interaction [24]. This suggests that the enhanced HPV-retromer association in cells depleted of Rab9a blocks HPV exit from the endosome independent of the effect of Rab9a knockdown on Rab7. Because both expression of DN Rab7 or depletion of Rab9a results in HPV accumulation in endosomes, however, it is difficult to uncouple actions of Rab9a and Rab7 on endosome exit. Nevertheless, the fact that Rab9a interacts with VPS35 [38] and Rab9a knockdown increases HPV-retromer association in cells expressing DN Rab7 is consistent with the possibility that Rab9a interferes with the HPV-retromer association independently of Rab7 and suggests that Rab9a antagonizes a Rab7-independent retromer recruitment mechanism. For example, it is possible that retromer is recruited to the HPV-containing endosomal vesicle via sorting nexins (SNXs) such as SNX3 independently of Rab7. In support of this notion, *in vitro* studies showed that the addition of SNX3 increased HPV16 L2 interaction with retromer [40] and that SNX3 can recruit retromer to liposomes in the absence of Rab7 or in the presence of DN Rab7 (albeit to a much lesser extent compared to recruitment in the presence of both SNX3 and CA Rab7) [41]. It is also possible that Rab9a knockdown may cause HPV accumulation in endosomes via both Rab7-independent and Rab7-dependent pathways.

A previous report claimed that DN Rab9a inhibits HPV infection [15], in contrast to our findings. Day et al. [15] used DsRed-tagged DN or wild-type Rab9a expressed in 293TT cells [15] whereas we assessed the effects of HA-tagged CA Rab9a. Furthermore, we measured HPV infectivity in the same cell population (containing cells that expressed CA Rab9a and those that did not), which showed that HPV infectivity is decreased by excess GTP-Rab9a (i.e., in cells expressing CA Rab9a). Finally, as an independent approach to increase GDP-Rab9a, DENDD2 knockdown HeLa cells displayed increased HPV infectivity compared to control cells (Fig. 5D and S6). These multiple lines of evidence support our conclusion that excess GDP-Rab9a promotes HPV infection whereas excess GTP-Rab9a impairs it. Nevertheless, these differences raise the possibility that various forms of Rab9a may support HPV infection in different ways depending on subtle differences in assay conditions.

Our findings also reveal that HPV and cellular protein cargos employ host trafficking machinery in different ways. Excess of the canonically active GTP-Rab9a and GTP-Rab7 support retrograde trafficking of cellular proteins [24, 36, 37] (Fig. 6), whereas excess of the canonically inactive GDP-Rab9a or the cycling between GTP- and GDP-bound Rab7 support HPV entry. Thus, although Rab9a and Rab7 are required for both HPV entry and cellular protein trafficking, cellular cargo utilizes these trafficking proteins in distinct ways compared to HPV. GDP-bound-Rab-supported HPV entry appears to differ from other viruses which are reported to be dependent on GTP-bound Rab proteins [4, 42, 43]. Although the trafficking of the Hepatitis C Virus (HCV) is mediated by GTP-bound Rab proteins [43], the assembly of HCV requires GDP-bound Rab32 [44]. Thus, it is possible that viruses may employ host factors in a distinct way. These observations indicate that non-canonical roles of Rab proteins or other host proteins may support virus infection.

If the cellular cargo trafficking machinery is limiting and if HPV uses these Rab proteins in the same way as the cellular protein cargo, the HPV entry process might compete with cellular protein trafficking and interfere with this process and trigger a host response that would inhibit virus infection. Alternatively, if cellular cargo out-competes the virus, the virus entry would be impaired. Thus, HPV may have evolved to use the cellular trafficking system in a distinct way from cellular protein cargo to avoid competition for the entry machinery and ensure successful entry. Those distinct trafficking strategies may also allow the development of therapeutic approaches to control HPV infection. For example, it may be possible to trap Rab9a in its GDP-bound form to block HPV entry while retaining cellular cargo trafficking.

Our findings also suggest a potential link between lipid metabolism and HPV infection, specifically that cholesterol accumulation may affect HPV infection by modulating Rab9a. Cholesterol accumulation in Niemann-Pick type C (NPC) 1-deficient cells reduces Rab9 extraction from cell membrane by GDI (guanine nucleotide dissociation inhibitor) [45]. The sequestered Rab9 appears to be GDP-bound form in these cells because the defective transport of cation-dependent mannose 6-phosphate receptor (CD-MPR) can be restored by overexpression of wild-type Rab9, but not by overexpression of DN Rab9 [45]. Thus, when cholesterol levels are high, there is likely less GTP-Rab9 and more GDP-Rab9a, which would support HPV entry.

In summary, we establish that Rab9a plays an essential role in HPV trafficking. Rab9a facilitates HPV exit from endosomes during entry by regulating retromer association to HPV. Our data also suggest that different Rab proteins operate at different times and in different ways during HPV entry, with HPV being handed over from early-acting Rab9a to later-acting Rab7. Our findings also uncover an unusual role of GTP- and GDP-bound forms of Rab9a in HPV trafficking. The inhibition of HPV entry by excess GTP-Rab9a and its promotion by excess GDP-Rab9a is in contrast to most cellular cargos, which require GTP-bound Rab proteins but not GDP-bound ones [1, 3, 6, 36, 37].

## Materials and Methods

### Cell lines

HeLa S3 cells were purchased from American Type Culture Collection (ATCC). HaCaT cells were purchased from AddexBio Technologies. 293TT cells were generated by introducing SV40 Large T antigen cDNA into HEK293T cells to increase Large T antigen expression and obtained from Christopher Buck (NIH). All cell lines were cultured at 37°C and 5% CO_2_ in Dulbecco’s modified Eagle’s medium (DMEM) supplemented with 20 mM HEPES, 10% fetal bovine serum (FBS), L-glutamine, and 100 units/mL penicillin streptomycin (DMEM10).

### Production of HPV pseudovirus (PsV)

HPV16 PsVs were produced by co-transfecting 293TT cells with wild-type p16SheLL-3XFLAG tag [14] together with pCAG-HcRed (Addgene #11152) or pCINeo-GFP [obtained from C. Buck] using polyethyleneimine (MilliporeSigma). For HPV18 and HPV5, p18SheLL-HA tag and p5SheLL were used, respectively. PsVs were purified by density gradient centrifugation in OptiPrep (MilliporeSigma) as described [46]. For comparing wild-type and 3R mutant, p16SheLL-HA WT and 3R mutant [16] were used. Briefly, cells were washed with Dulbecco’s Phosphate Buffered Saline (DPBS) at 24 h post transfection, incubated in DMEM10, and collected 72h post transfection in siliconized tubes. The cells were then incubated in lysis buffer (DPBS with 0.5% Triton X-100, 10 mM MgCl_2_, 5 mM CaCl_2_, 100 µg/mL RNAse A (Qiagen)) overnight at 37°C in a water bath to allow capsid maturation. The lysates containing matured PsVs were loaded on an OptiPrep gradient that had been stabilized at least 1 h and centrifuged at 50k x g for 3.5-4 h at 4°C in a SW-55Ti rotor (Beckman). Fractions were collected in siliconized tubes and subjected to SDS-PAGE followed by Coomassie blue staining to assess the abundance of L1 and L2 proteins. Peak fractions were pooled, aliquoted, and stored at -80°C.

### Determining HPV PsV infectivity

0.37×10^5^ HeLa S3 cells per well were plated in 24-well plates ∼48 h prior to infection. Approximately 6 h later, cells were transfected with 6.7 nM of indicated siRNAs (Table S1) using Lipofectamine RNAiMAX (Invitrogen) according to manufacturer’s protocol. Non-targeting siRNA (Table S1) was used as a negative control. 40-48 h after transfection, cells were infected with PsVs at an infectious MOI of ∼5 or ∼0.5 in unmodified HeLa cells. At 48 hpi, infectivity was determined by using flow cytometry to measure fluorescence intensity produced from expression of the reporter gene. Relative percent infectivity was determined by normalizing mean fluorescence intensity of samples transfected with experimental siRNA to that of the cells transfected with control siRNA, which was set at 100%.

### Western blot analysis

siRNA-treated cells were lysed using ice-cold 1X Radioimmunoprecipitation assay (RIPA) [50 mM Tris (pH 7.4), 150 mM NaCl, 1% Nonidet P-40, 1% sodium deoxycholate, 0.1% sodium dodecyl sulfate, 1 mM Ethylenediaminetetraacetic acid] buffer supplemented with 1X HALT protease inhibitor cocktail (Pierce) for 15 min at 4°C. After centrifugation at 14,000 rpm for 15 min at 4°C, the protein concentration in the supernatant was determined by Bicinchoninic acid (BCA) protein assay (Pierce). After normalization for protein amounts, the supernatant was mixed with 4X Laemmli dye (Bio-rad) supplemented with 10% 2-mercaptoethanol and incubated in a water bath for 7 min at 100°C. Samples were then separated by SDS-PAGE (4-12% gel) (Bio-rad) and analyzed by Western blotting using antibodies recognizing Rab9a (Thermo Fisher, 11420-1-AP, 1:1,000), Rab7 (Cell Signaling Technology, 9367, 1:1,000), TBC1D5 (Abcam, ab203896, 1:1,000), pan actin (Cell Signaling Technology, 4968, 1:2,000), FLAG (Sigma, A8592, 1:1,000) or GST (Santa Cruz, sc-138, 1:2,000). Secondary horseradish peroxidase (HRP)-conjugated antisera recognizing rabbit or mouse antibodies as appropriate (Jackson ImmunoResearch, 711-035-152, 115-035-146) were used at 1:5,000 dilution. The blots were developed with SuperSignal West Pico or Femto Chemiluminescent substrate (Pierce) and were visualized by using a FluorChem Imager (Bio-technne, FE0685) or film processor (Fujifilm).

### Proximity ligation assay (PLA)

0.35×10^5^ HeLa S3 cells per well were plated in 24-well plates containing glass coverslips 48 h prior to infection. Approximately 6 h later, cells were transfected with 6.7 nM of indicated siRNA (Table S1) as described above. At 40-48 h after transfection, cells were infected with PsVs at MOI of ∼200 in unmodified HeLa cells. As indicated, DMSO (0.2% as final concentration), 100 nM BafA, or 2 µM XXI were added to the medium 30 min prior to infection. At indicated times post-infection, cells were fixed with 4% paraformaldehyde (Electron Microscopy Sciences) at room temperature (RT) for 12 min, permeabilized with 1% Saponin (Sigma-Aldrich) at RT for 35-40 min, and blocked using DMEM10 at RT for 1-1.5 h. Cells were then incubated overnight at 4°C with a pair of mouse and rabbit antibodies; a mouse antibody recognizing L1 (BD Biosciences, 554171, 1:1,000 when used with anti-TGN46, 1:200 when used with other antibodies) or mouse antibodies recognizing cellular proteins (anti-EEA1, BD Biosciences, 610457, 1:100; anti-VPS35, Abcam, ab57632, 1:200); rabbit antibodies recognizing cellular proteins (anti-TGN46, Abcam, ab50595, 1:600; anti-EEA1, Cell Signaling Technology, 2411, 1:75; anti-VPS35, Abcam, ab157220, 1:200). PLA was performed with Duolink reagents (Sigma Aldrich) according to the manufacturer’s instructions as described [47]. Briefly, cells were incubated in a humidified chamber at 37°C with a pair of PLA antibody (mouse and rabbit) probes for 75 min, with ligation mixture for 45 min, and then with amplification mixture for 3 h, followed by series of washes. Nuclei were stained with 4,6-diamidino-2-phenylindole (DAPI). Cellular fluorescence was imaged using the Zeiss LSM980 confocal microscope. Images were processed using a Zeiss Zen software version 3.1 and quantified using Image J Fiji version 2.3.0/1.53f.

### Co-immunoprecipitation of Rab9a and L2

7×10^5^ HeLa S3 cells per pate were plated in 6 cm^2^ plates then infected with PsVs at the MOI of ∼100. As indicated, 100 nM BafA was added 30 min prior to infection (non-treated control cells were treated with DMSO alone (0.1-0.2% as final concentration)). At 6.5 hpi, cells were washed twice with DPBS and cross-linked with 1.5 mM DSP in DPBS for 30 min at RT. The reaction was then quenched with 100 mM Tris HCl (pH 7.4) for 15 min at RT. The cells were then washed three times with ice cold DPBS and lysed in 500 µL of lysis buffer (20 mM HEPES [pH 7.4], 50 mM NaCl, 5 mM MgCl_2_, 1% *n*-Dodecyl β-D-maltoside (DDM) and 0.05% Triton X-100) supplemented with 1 X HALT protease and phosphatase inhibitor cocktail (Thermo Fisher). The lysate was centrifuged at 14,000 rpm for 20 min at 4°C and supernatant was transferred to new tubes and normalized for total protein amounts determined by BCA assay. 50 µL of supernatant was reserved for input samples. The remainder of the supernatant was incubated with 6 µl of anti-Rab9a antibodies overnight at 4°C, then incubated with 40 µl of protein G magnetic beads (Thermo Fisher Scientific) for one hour at room temperature. Bound proteins were collected with magnet, washed three times with the lysis buffer, and eluted with 50 µL of 2X Laemmli sample buffer (Bio-Rad) at 95°C, followed by SDS-PAGE and Western blot analysis carried out as described above.

### RILP pull down assay

Plasmids expressing GST-RILP (obtained from Christopher Burd, Yale University) or GST alone (from pGEX KG vector; Addgene #77103) were transformed into *Escherichia coli* strain BL21(DE3). Bacterial cultures at OD_600_ of 0.5-0.6 were induced by 0.5 mM isopropyl β-D-1-thiogalactopyranoside (IPTG) at 22°C for 5 h. Bacteria were harvested, washed with Tris-Buffered Saline (TBS) (50 mM Tris [pH 8.0], 150 mM NaCl), and lysed with B-per lysis buffer (Thermo Scientific) supplemented with 1X HALT protease inhibitor cocktail (Pierce). GST-RILP or GST proteins were purified by using a pre-equilibrated slurry of glutathione beads (Thermo Scientific, #16100) in TBS containing 1mM MgCl_2_ and washed three times in the same buffer containing 0.05% Triton X-100. Purified proteins were eluted from the beads with reduced 20mM glutathione in TBS containing 1mM MgCl_2_ and dialyzed into HEPES buffer (20 mM HEPES [pH 7.4], 50 mM NaCl, 5 mM MgCl_2_, 1 mM dithiothreitol (DTT)) using Slide-A-Lyzer dialysis Cassette. Protein amounts were quantified with bovine serum albumin (BSA) standards separated by SDS-PAGE followed by Coomassie blue staining.

2×10^5^ HeLa S3 cells per plate were plated in 6 cm^2^ plates and transfected with 6.7 nM of indicated siRNAs (Table S1) as described above. 40-48 h after transfection, cells were infected with PsVs at MOI of ∼150 in unmodified HeLa cells. At 12 hpi, cells were lysed using 400 µL ice-cold lysis buffer (HEPES buffer containing 0.15% Triton X-100) supplemented with 1X HALT protease and phosphatase inhibitor cocktail. After centrifugation at 14,000 rpm for 20 min at 4°C, the protein concentration in the supernatant was determined by BCA protein assay (Pierce). After normalization for protein amounts, the supernatant was incubated with 15 µg of purified GST or GST-RILP proteins at 4°C for 2 h in 500 µl of lysis buffer. 40 µL of mixtures were taken for input samples. Then 40 µL of pre-equilibrated (using lysis buffer) slurry of glutathione beads were added and the mixtures were further incubated at 4°C for 3 h, followed by three washes in lysis buffer. Bound proteins were eluted with 50 µL of 2X Laemmli sample buffer containing 5% 2-mercaptoethanol at 95°C for 5 min. Input and eluted samples were separated by SDS-PAGE, and Western blot analysis was carried out as described above.

### Construction of plasmids

Plasmids expressing HA-tagged Rab9a CA or DN variants were constructed as follows: CA and DN mutant *Rab9a* genes were amplified from JB84 (Addgene #128908) and DsRed-rab9 DN (Addgene, Plasmid #12676), respectively, using primers HA-Rab9a-BamHI-F. (TGA ACC GTC AGA TCG CCT GGA GAA GGA TCC ATG TAC CCA TAC GAC GTT CCA GAT TAC GCT GCA GGA AAA TCA TCA CTT TTT AAA GTA) and Rab9a-EcoRI-R (GAA AAG CGC CTC CCC TAC CCG GTA GAA TTC TCA ACA GCA AGA TGA GCT AGG CTT GGG CTT), then introduced between BamHI and EcoRI sites of pRetroX-Tight-Pur vector (Takara, 632105). Resulting plasmids containing the desired mutation were confirmed by DNA sequencing.

### Determining HPV PsV infectivity in cells transiently expressing Rab9a CA

0.35×10^5^ 293TT cells per plate were plated in 24-well plates 16-20 h prior to transfection of plasmids encoding HA-tagged CA Rab9a or the empty vector as control (Takara, 632105). 48 h after transfection, cells were infected with PsVs at MOI of ∼1 in unmodified HeLa cells. 48 hpi, cells were fixed with 4% paraformaldehyde, permeabilized with 1% Saponin, and blocked with 3% BSA. Cells were then stained using PE-conjugated antibodies recognizing HA (MACS Molecular, 130-120-717, 1:1000), followed by three to four times of washes using DPBS containing 0.1% Tween-20. HA-tagged Rab9a CA or DN protein abundance and HPV PsV infectivity were determined by flow cytometry measuring fluorescence intensity produced by PE-labeled proteins and expressed GFP simultaneously.

### Cellular protein DMT1-II trafficking assay

To analyze cells overexpressing Rab9a CA, 0.3×10^5^ 293TT cells per well were plated in 24-well plates containing glass coverslips 16-20 h prior to transfection of 2 µg plasmid encoding HA-tagged CA Rab9a, followed by transfection of 1 µg plasmid encoding DMT1-II-GFP the next day. 24 h after the second transfection, cells were processed as described below. To examine cells depleted for DENND2D, 0.3×10^5^ 293TT cells per plate were plated in 24-well plate containing glass coverslips and transfected with 6.7 nM of indicated siRNAs (Table S1) as described above. 40-48 h post siRNA-transfection, cells were then transfected with 1 µg plasmid encoding DMT1-II-GFP. 24 h after the second transfection, samples were fixed with 4% paraformaldehyde at RT for 10 min, permeabilized with 1% Saponin at RT for 35-40 min, and blocked using DMEM10 at RT for 1-1.5 h. Cells were then incubated overnight at 4°C with a mouse antibody recognizing GFP (Santa Cruz, sc-9996, 1:300) and a rabbit antibody recognizing TGN46 (Abcam, ab50595, 1:250). A rat antibody recognizing HA was used (Roche, 11867423001, 1:250) to visualize HA-tagged CA Rab9a. Samples were then stained with 1:200 Alexa-Fluor-conjugated secondary antibodies (Life Technologies) at RT for one hour. Nuclei were stained with DAPI. Cells were imaged using the Zeiss LSM980 or LSM880 confocal microscope. Images were processed as described above in PLA section.

### Statistical analyses

For comparisons of two groups, unpaired *t*-tests were applied. For comparisons of more than three groups, One-way ANOVA with the ordinary ANOVA test. These analyses provide *P*-values for each comparison.

## Acknowledgments

We would like to thank Qun Lin for making HPV18 and HPV5 PsVs. We also thank Changin Oh for providing plasmid DNAs of p16SheLL-HA WT and 3R mutant. We also thank Yuka Takeo and Jian Xie for their technical advice for performing proximity ligation assay and co-immunoprecipitation analysis. Flow Cytometry was conducted in the flow cytometry shared resource of the Yale Cancer Center. This work was supported by grants from the NIH to DD (R35-CA242462).

## Figure Legends

**Fig. S1.**
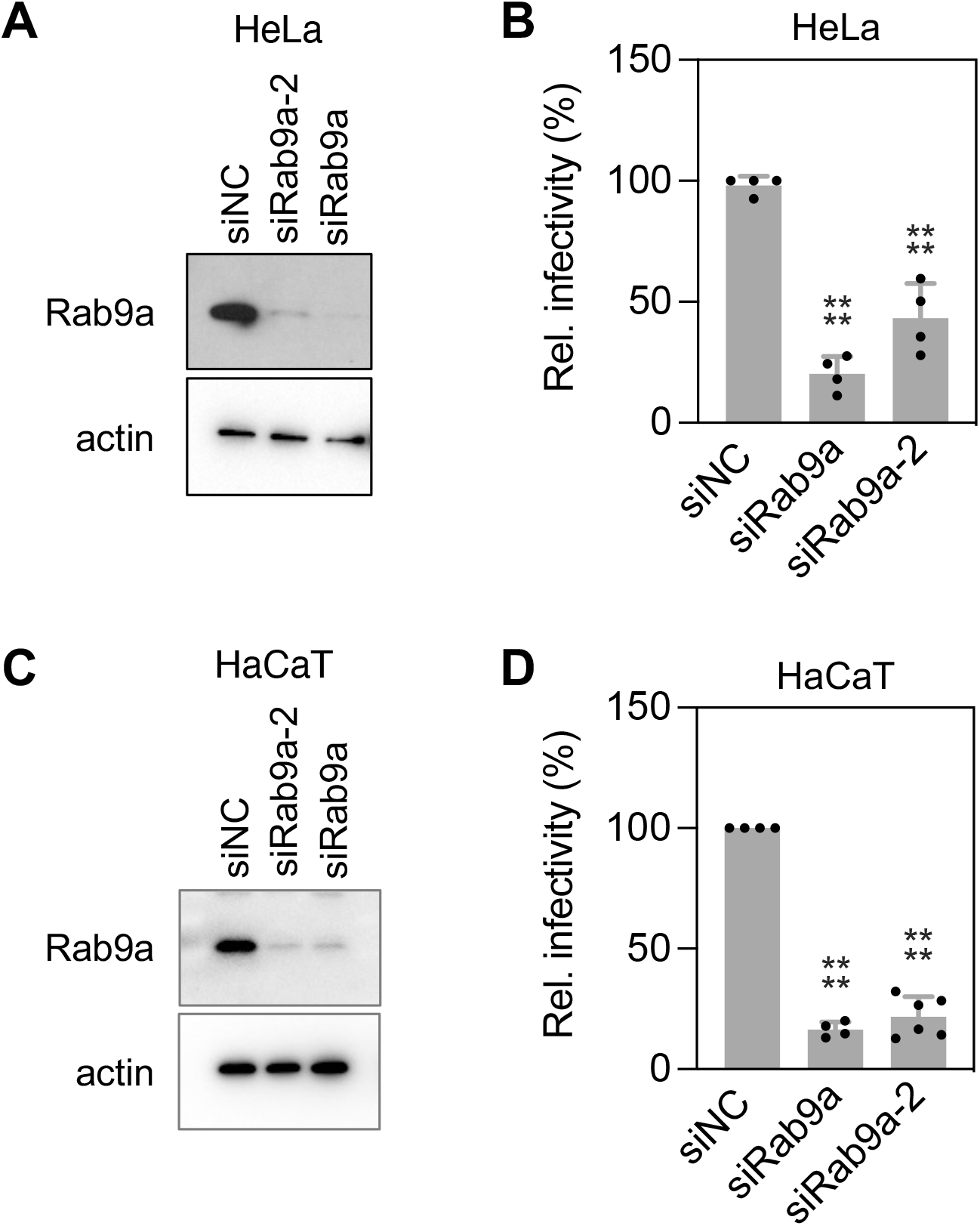
Rab9a knockdown inhibits HPV infection in both HeLa and HaCaT cells. (**A**) HeLa S3 cells were transfected with siNC or two different siRNAs targeting Rab9a (siRab9a and siRab9a-2) and were subjected to Western blot analysis using antibodies recognizing Rab9a and actin as a loading control. (**B**) siRNA-treated cells as described in (**A**) were mock-infected or infected at the MOI of ∼2 with HPV16 PsV L2-3XFLAG containing the GFP reporter plasmid. At 48 hpi, GFP fluorescence was determined by flow cytometry. The results are shown as percent relative infectivity (based on mean fluorescence intensity) normalized to siNC treated cells (*right*). Each dot shows the result of an individual experiment. ****, *P* < 0.0001. (**C**) As in (**A**) except using HaCaT cells. (**D**) As in (**B**) except using HaCaT cells.

**Fig. S2.**
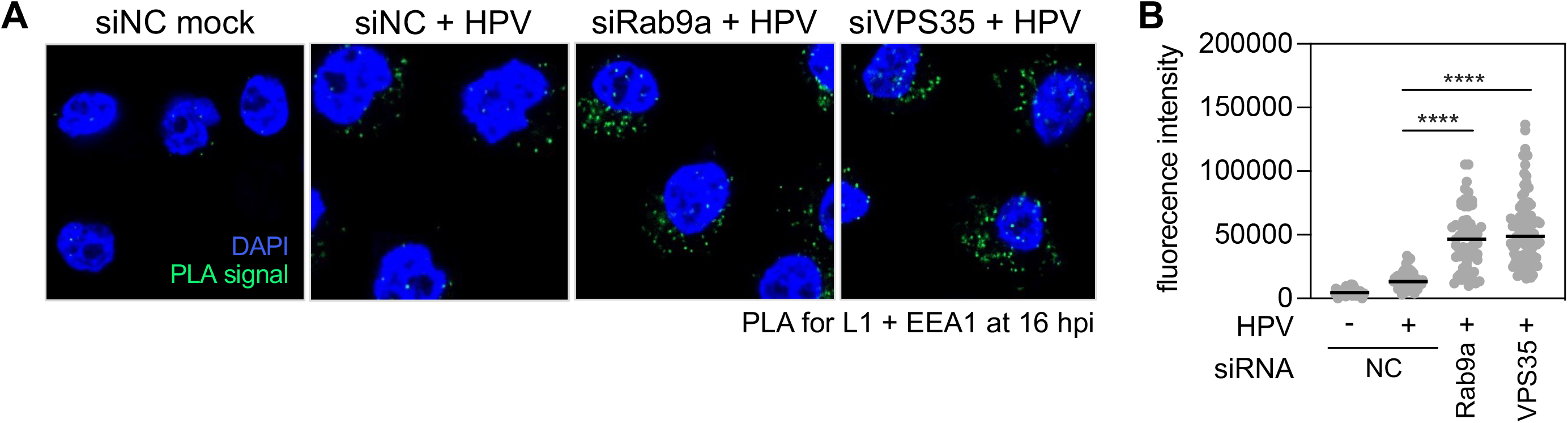
Knockdown of Rab9a or VPS35 causes HPV accumulation in endosomes. (**A**) HeLa S3 cells were transfected with siNC, siRab9a or siVPS35 and mock-infected or infected at the MOI of ∼200 with HPV16 PsV L2-3XFLAG containing the HcRed reporter plasmid. At 16 hpi, PLA was performed with antibodies recognizing HPV L1 and EEA1. Mock, uninfected; HPV, infected. PLA signals are green; nuclei are blue (DAPI). Similar results were obtained in two independent experiments. (**B**) The fluorescence of PLA signals was determined from multiple images obtained as in (**A**). Each dot represents an individual cell (*n*>40) and black horizontal lines indicate the mean value of the analyzed population in each group. ****, *P* < 0.0001. The graph shows results of a representative experiment. Similar results were obtained in two independent experiments.

**Fig. S3.**
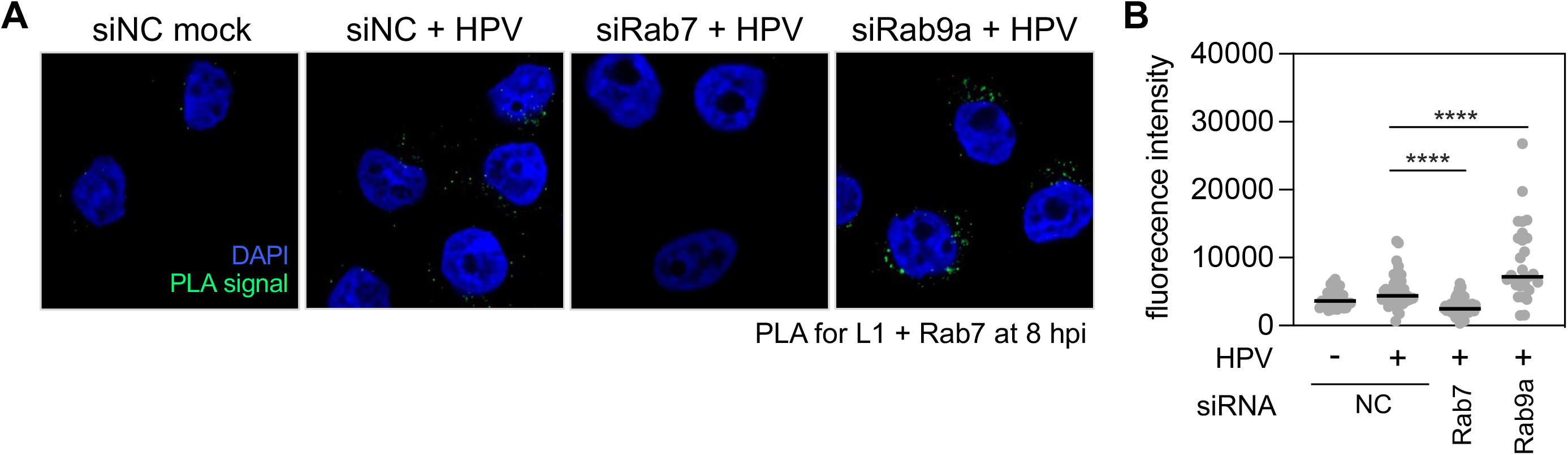
Rab9a knockdown increases Rab7 association to HPV. (**A**) HeLa S3 cells were transfected with negative control siNC, siRab7, or siRab9a and mock-infected or infected at the MOI of ∼200 with HPV16 PsV L2-3XFLAG containing the HcRed reporter plasmid. At 8 hpi, PLA was performed with antibodies recognizing HPV L1 and Rab7. Mock, uninfected; HPV, infected. PLA signals are green; nuclei are blue (DAPI). Similar results were obtained in two independent experiments. (**B**) The fluorescence of PLA signals was determined from multiple images obtained as in (**A**). Each dot represents an individual cell (*n*>25) and black horizontal lines indicate the mean value of the analyzed population in each group. ****, *P* < 0.0001. The graph shows results of a representative experiment. Similar results were obtained in two independent experiments.

**Fig. S4.**
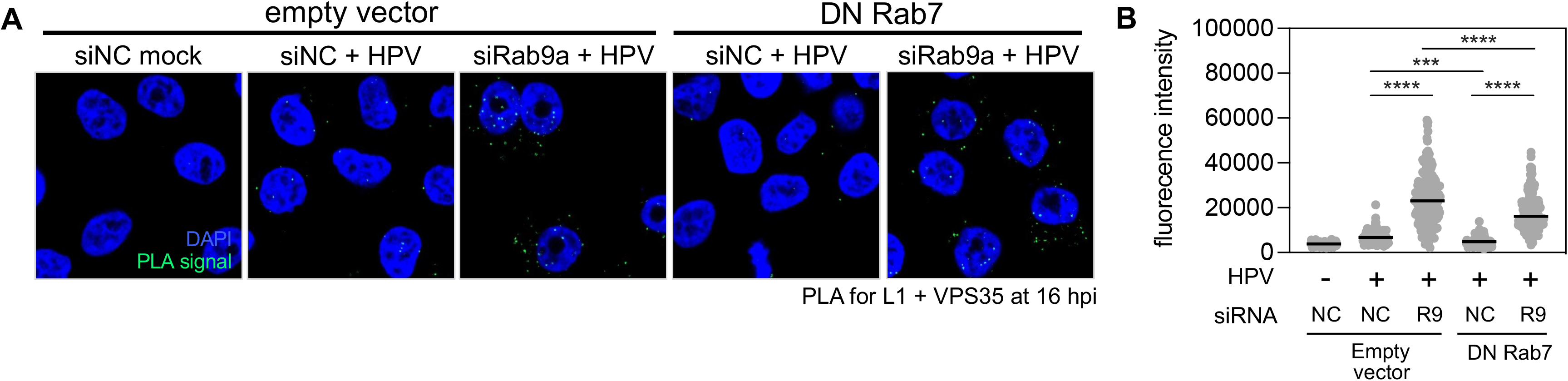
Rab9a knockdown increases HPV-retromer association even in cells overexpressing GDP-Rab7. (**A**) HeLa S3 cells stably transduced with an empty vector or a plasmid expressing dominant negative Rab7 (Rab7 DN) were transfected with siNC or siRab9a, and mock-infected or infected at the MOI of ∼200 with HPV16 PsV L2-3XFLAG containing the HcRed reporter plasmid. At 16 hpi, PLA was performed with antibodies recognizing HPV L1 and VPS35. PLA signals are green; nuclei are blue (DAPI). Similar results were obtained in two independent experiments. (**B**) The fluorescence of PLA signals was determined from multiple images obtained as in (**A**). Each dot represents an individual cell (*n*>30) and black horizontal lines indicate the mean value of the analyzed population in each group. ***, *P* < 0.001; ****, *P* < 0.0001. The graph shows results of a representative experiment. Similar results were obtained in two independent experiments.

**Fig. S5.**
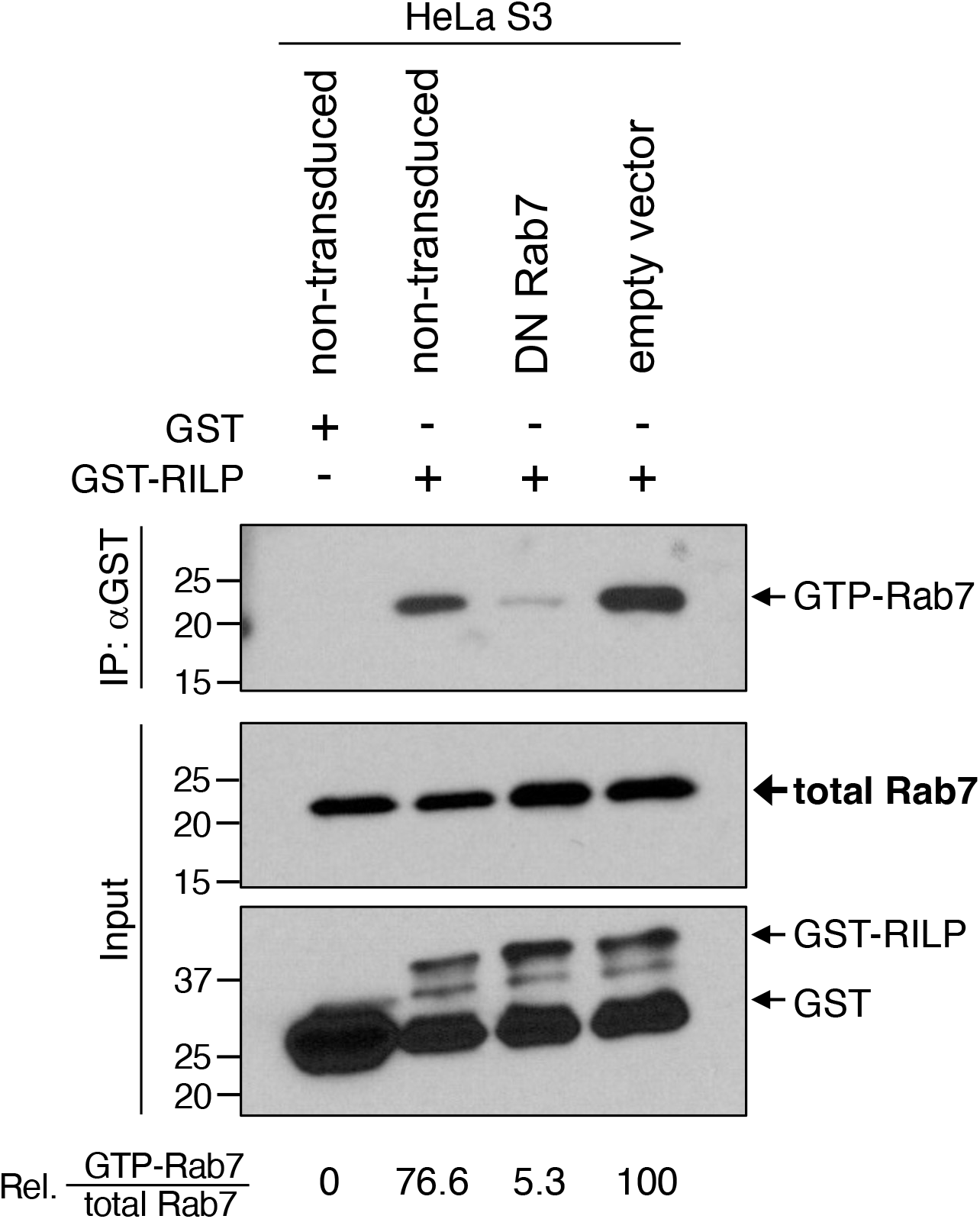
Expressing DN Rab7 depletes most GTP-Rab7. Lysates from HeLa S3 cells or those stably transduced with an empty vector or a plasmid expressing dominant negative Rab7 (Rab7 DN) were pulled down with GST or GST-RILP (IP) and subjected to Western blot analysis using antibodies recognizing Rab7 or GST together with samples not pulled down (input). Numbers below the bottom panel indicate relative percent abundance of GTP-Rab7 normalized to cells transfected with empty vector (set at 100%).

**Fig. S6.**
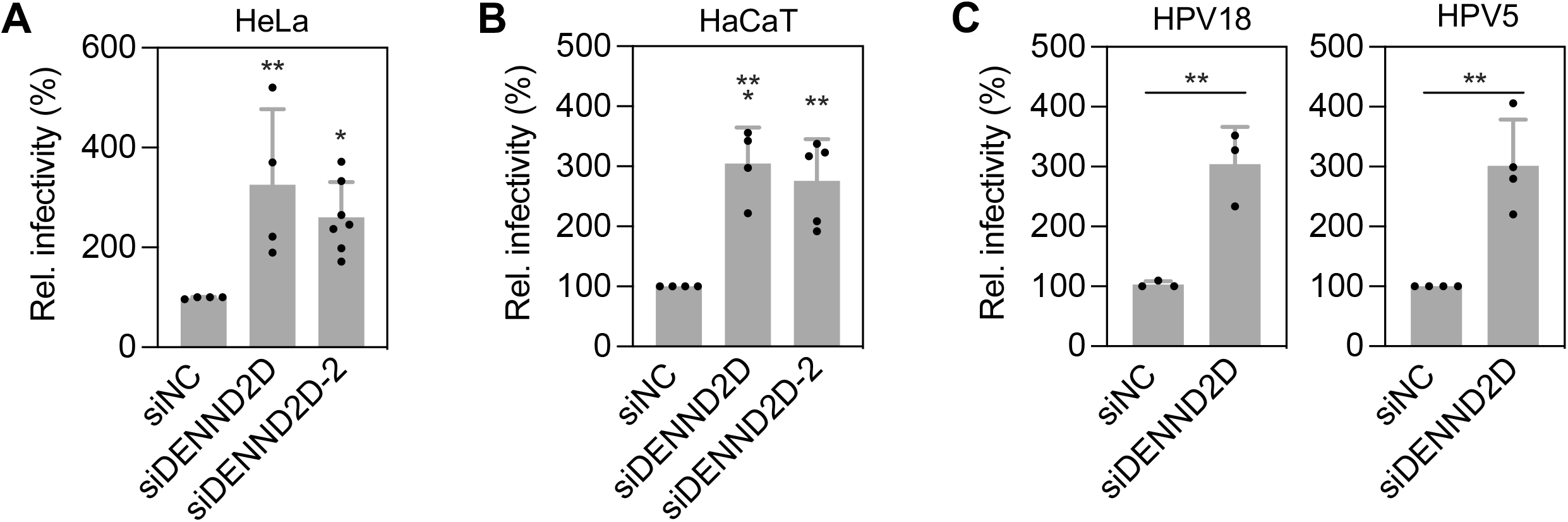
DENND2D knockdown increases HPV infection in both HeLa and HaCaT cells. (**A**) HeLa S3 cells were transfected with siNC or two different siRNAs targeting DENND2D (siDENND2D and siDENND2D-2) and mock-infected or infected at the MOI of ∼0.2 with HPV16 PsV L2-3XFLAG containing the GFP reporter plasmid. At 48 hpi, GFP fluorescence was determined by flow cytometry. The results are shown as percent relative infectivity (based on mean fluorescence intensity) normalized to the siNC treated cells (set at 100%). Each dot shows the result of an individual experiment. *, *P* < 0.05; **, *P* < 0.01. (**B**) As in (**A**) except using HaCaT cells. ***, *P* < 0.001. (**C**) As in (**A**) except cells were infected with HPV18 and HPV5.

**Table S1.**
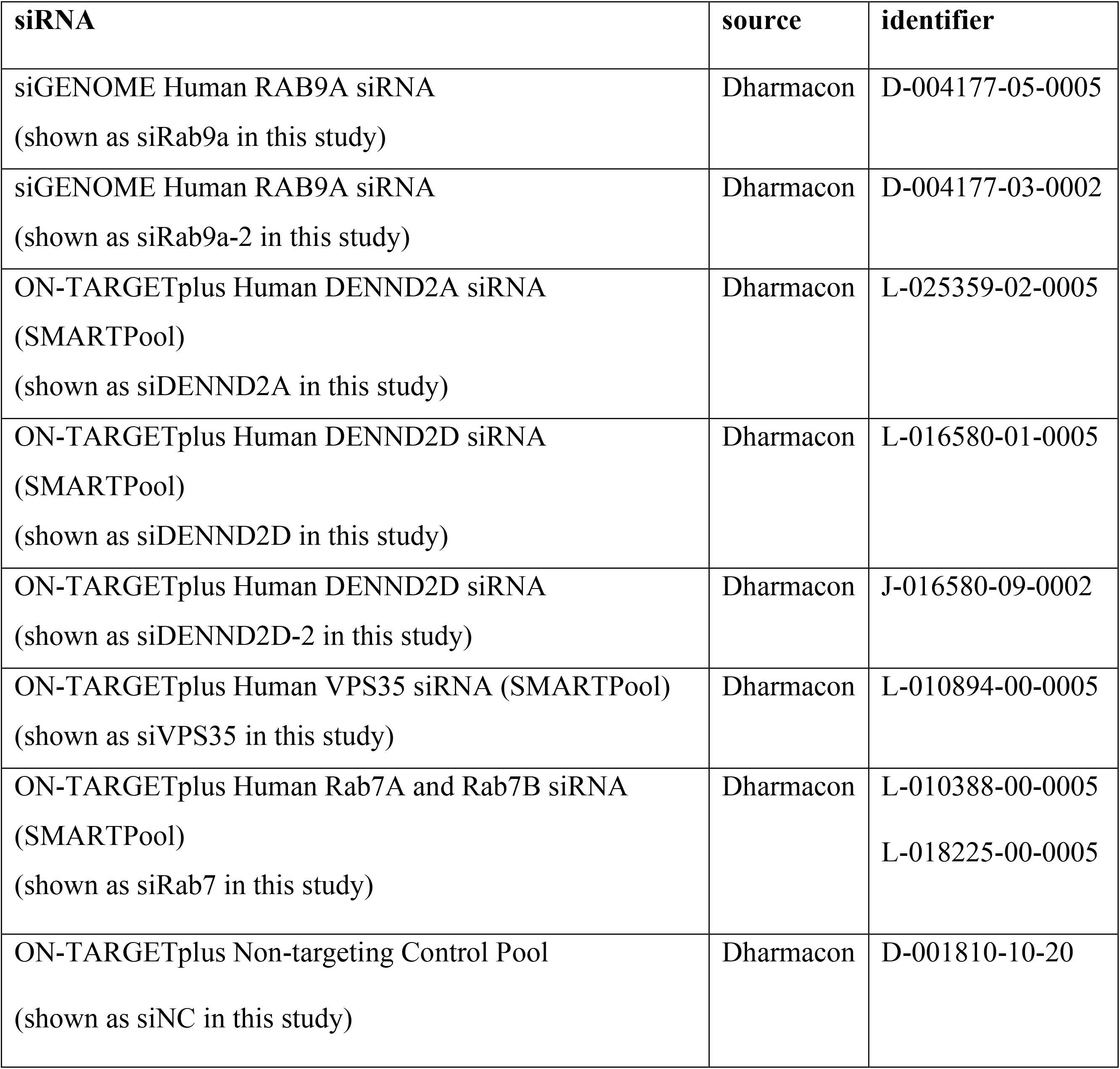
Oligonucleotides used in this study.

| **siRNA** | **source** | **identifier** |
| --- | --- | --- |
| siGENOME Human RAB9A siRNA<br>(shown as siRab9a in this study) | Dharmacon | D-004177-05-0005 |
| siGENOME Human RAB9A siRNA<br>(shown as siRab9a-2 in this study) | Dharmacon | D-004177-03-0002 |
| ON-TARGETplus Human DENND2A siRNA<br>(SMARTPool)<br>(shown as siDENND2A in this study) | Dharmacon | L-025359-02-0005 |
| ON-TARGETplus Human DENND2D siRNA<br>(SMARTPool)<br>(shown as siDENND2D in this study) | Dharmacon | L-016580-01-0005 |
| ON-TARGETplus Human DENND2D siRNA<br>(shown as siDENND2D-2 in this study) | Dharmacon | J-016580-09-0002 |
| ON-TARGETplus Human VPS35 siRNA (SMARTPool)<br>(shown as siVPS35 in this study) | Dharmacon | L-010894-00-0005 |
| ON-TARGETplus Human Rab7A and Rab7B siRNA<br>(SMARTPool)<br>(shown as siRab7 in this study) | Dharmacon | L-010388-00-0005<br>L-018225-00-0005 |
| ON-TARGETplus Non-targeting Control Pool<br>(shown as siNC in this study) | Dharmacon | D-001810-10-20 |

## Notes

### Competing Interest Statement

The authors have declared no competing interest.

